# Microbiome-host interactions drive a trade-off between sleep quality and lifespan in *Drosophila*

**DOI:** 10.64898/2025.12.24.696379

**Authors:** Priyanshu Bhargava, Shenyue Chen, Jinho Lee, Yuto Yoshinari, Takashi Nishimura, Masayuki Ushio, Yuki Sugiura, Kwun For Cheng, Yu Shan Fung, Koji Hase, Hugh Nakamura, Masato Enomoto, Jung Kim, Heath E. Johnson, Yi Liao, Yukinori Hirano

## Abstract

Understanding the interactions between various aging processes and the resulting heterogeneity in aging is crucial for promoting healthy aging. Here, we provide evidence that heterogeneity in microbiome and host interactions contributes to diversifying aging phenotypes in sleep, gut integrity, and longevity in *Drosophila*. Aged flies exhibiting sleep fragmentation preserve gut integrity, accompanied by a shift in microbiota composition, particularly an increase in *Acinetobacter junii*. *A. junii* induces sleep fragmentation via its metabolite, urocanic acid, through serotonin receptor-dependent dopamine upregulation. In parallel, *A. junii* exploits the host response to promote its growth, leading to lifespan extension, which is recapitulated by genetically modified *Escherichia coli*, suggesting a trade-off between sleep quality and lifespan. Our study demonstrates a systematic mechanism underlying aging heterogeneity, suggesting interventions through bacterial supplements.

## Introduction

Although aging is inevitable for all living organisms, the onset and degree of age-related dysfunctions differ between individuals (*1*). In addition to genetic factors, internal or external variables likely contribute to generating heterogeneity in age-related phenotypes. Even laboratory model animals carrying relatively similar genetic backgrounds reproducibly exhibit heterogeneity in aging phenotypes; for example, some die earlier, while others live longer. The factors inducing heterogeneity in aging may be common or different across cells and organs, or they could be shaped by communication among these entities. Furthermore, the heterogeneity factors could be linked to interactions with the microbiome and host responses. By studying the variability that determines aging heterogeneity, the inevitable aging processes could become intervenable.

Aged animals, including humans, exhibit difficulties in initiating and maintaining sleep (*2*), which are hallmarks of insomnia. In humans, non-rapid eye movement (NREM) sleep is susceptible to impairment due to aging (*3, 4*), affecting approximately half of the population studied (*1*). Laboratory animals also exhibit age-dependent sleep impairment. Aged mice display fragmented NREM sleep, which is related to hyperactivity of hypocretin/orexin neurons in the lateral hypothalamus (*5*), although how this age-dependent change occurs is unknown. Flies, which sleep during the day and night while being more active at dawn and dusk, also demonstrate disruption of sleep homeostasis upon aging (*6*). Similar to the observation that human males display more pronounced impairment in NREM sleep than females (*7*), aged male flies exhibit sleep fragmentation (SF), defined as the breakdown of consolidated sleep into shorter, interrupted bouts, whereas females do not (*6*). SF in male flies can be alleviated by reducing insulin or target of rapamycin (TOR) signaling (*8*), which are considered general factors related to aging (*9*). However, the causal factors inducing age-dependent SF are currently unknown. As sleep is one of the major homeostatic controls coordinating both brain and body functioning, its dysfunction increases risks of morbidity and mortality, and negatively affects cognitive functions (*2, 10, 11*). We postulated that sleep loss in aged males could be an entry point to study the mechanisms of aging heterogeneity linking multiple phenotypes, and therefore characterized individual age-related biological traits accompanied by SF.

## Results

### The homeostasis of sleep and gut integrity is inversely correlated in aged flies

To monitor sleep patterns, we designed a circular arena to video-record the flies, where food intake could be simultaneously tested (fig. S1A). As previously reported (*6*), in both nighttime (ZT13-23) and daytime (ZT1-11), males at 25-day old exhibited a decrease in total sleep amount (Fig. 1A and fig. S1, C and D) and an increase in sleep episodes (fig. S1, E and F), indicating reduced sleep duration per episode, which we refer to as sleep fragmentation (SF) (fig. S1, G and H). Aged females conversely showed an increase in total sleep amount (fig. S1, B to D), a decrease in sleep episodes (fig. S1, E and F), and an increase in sleep duration per episode (fig. S1, G and H), indicating that SF is a specific aging phenotype in males as previously reported (*6*). We hereafter focused on males and their nighttime sleep since SF in aged males was similar during both nighttime and daytime. SF developed at 15-day old and was sustained at later ages, while a subpopulation was exempt from the emergence of SF (Fig. 1B). To understand the aging phenotypes associated with SF, we categorized aged flies with sleep durations per episode below the median as SF flies, while the remaining flies were classified as non-sleep fragmented (NSF) flies. Compared with aged males with NSF, those with SF lived longer (Fig. 1C) and preserved negative geotaxis (Fig. 1D), suggesting an unexpected inverse correlation between sleep homeostasis and physical conditions. Food intake of aged SF and NSF males was comparable (fig. S1I), suggesting that SF is not related to food intake. We further conducted RNA-seq analyses using individual bodies (Fig. 1E) and heads (Fig. 3) to determine the differentially expressed genes (DEGs) associated with SF. The gene expression profiles between young and aged male bodies with SF or NSF showed segregation between groups (Fig. 1F). Comparison between aged male bodies with SF and NSF indicated 96 DEGs exhibiting more than 2-fold difference (Fig. 1G), which can be further classified into four types by comparing with young bodies (Fig. 1H): type A (1 gene) showing increased expression in flies with SF, type B (63 genes) showing reduced expression in flies with NSF, type C (4 genes) showing reduced expression in flies with SF, and type D (28 genes) showing increased expression in flies with NSF (Fig. 1I and table 1). Among the four types, type B genes were enriched within a cluster associated with serine-type endopeptidase activity and serine hydrolase activity (Fig. 1J), while genes in other types did not show functional enrichment by gene ontology analysis. Type B genes include trypsin- and chymotrypsin-like proteases such as Jon65Ai (Fig. 1I), which are predominantly expressed in the midgut, presumably to support food digestion. We speculated that aged males with SF may maintain gut homeostasis, but those with NSF may not.

**Fig. 1.**
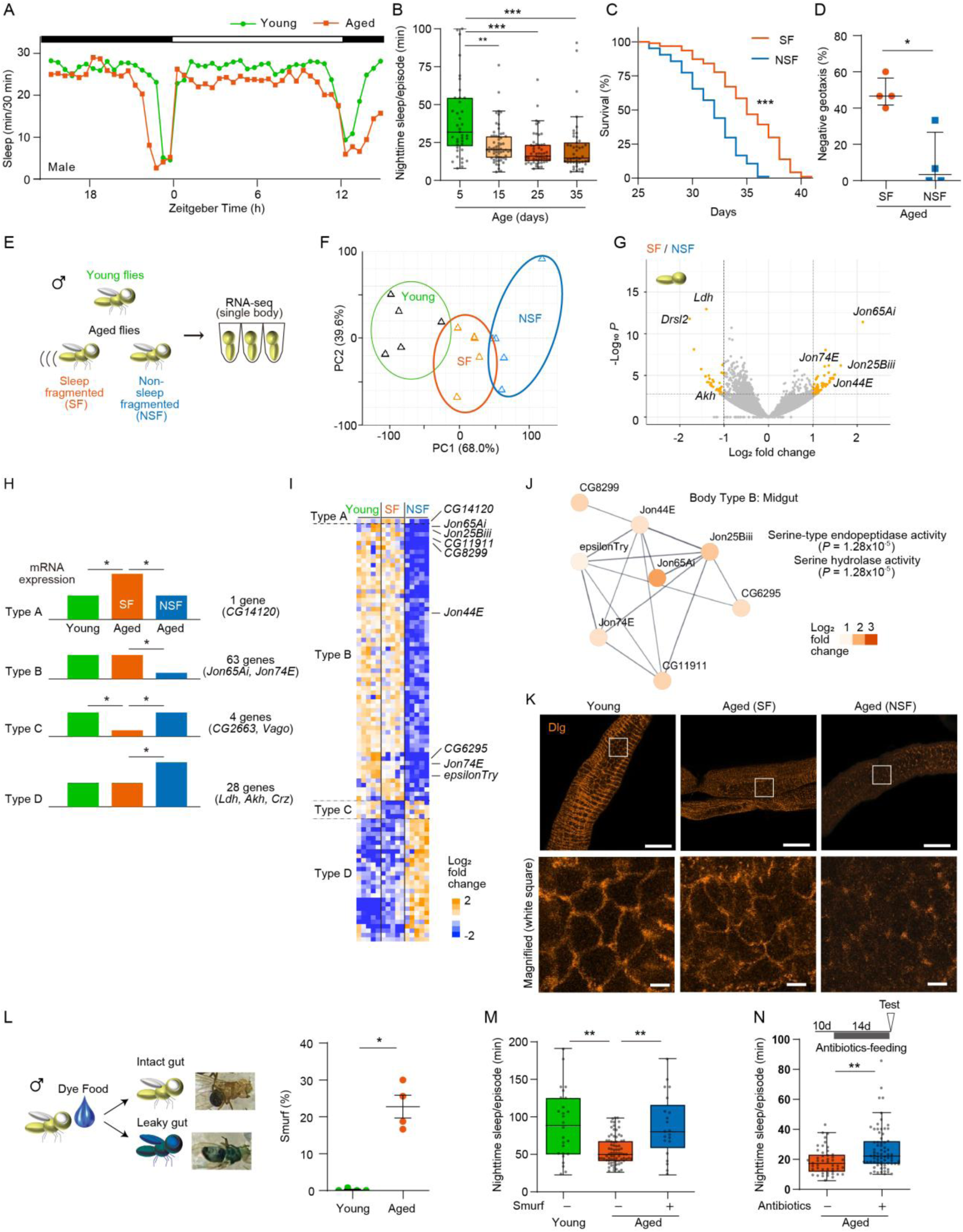
The homeostasis of sleep and gut integrity is inversely correlated in aged flies. **(A)** Sleep profile of males. The 5-day-old and 25-day-old flies were video-recorded to monitor the sleep duration every 30 min, and the averages were plotted against the Zeitgeber time (ZT). The light:dark cycle was indicated by white/black bars. Young, *n* = 41; aged, *n* = 53. **(B)** SF is developed after 15-day old. Sleep of the male flies at the indicated ages was analyzed. Kruskal-Wallis test, *P* < 0.0001; from left, *n* = 41, 62, 53, 46. **(C and D)** Survival (C) and negative geotaxis (D) of aged males with SF or NSF. Male flies showing SF or NSF at 25-day old were separated, and their survival or negative geotaxis were examined. (C) Log-rank test; SF, *n* = 94; NSF, *n* = 83. (D) Two-sided Mann-Whitney U-test; *n* = 4 for all. **(E)** Schematics of RNA-seq from individual fly bodies. **(F)** Principal component analysis (PCA) of mRNA expression data from the RNA-seq analysis. **(G)** Volcano plot representing the differentially expressed genes (DEG). **(H and I)** Classification (H), heatmap of the log_2_ fold changes in expression (I) of the DEGs. **(J)** Protein network within the type B genes identified from the STRING database. **(K)** Gut barrier is disrupted in aged males with NSF. Sleep was analyzed using male flies at 5-day or 25-day old, and gut was stained with anti-Dlg antibody. The images are representative of four experimental replicates. (upper) Scale bar: 50 μm. (lower) Scale bar: 5 μm. **(L)** Aged flies exhibit leaky gut. Groups of 5-day or 25-day-old flies were fed food containing blue dye for 1 day, and the flies showing blue dye in their body (Smurf flies) were counted. Two-sided Mann-Whitney U-test; *n* = 4. **(M)** Aged males showing intact gut display SF. Sleep was analyzed using flies at 5-day or 25-day old, while the flies were fed blue dye. Kruskal-Wallis test, *P* < 0.0001; from left, *n* = 26, 78, 21. **(N)** Feeding flies antibiotics suppresses SF. Flies were fed antibiotics as indicated, and sleep was analyzed using flies at 5-day or 25-day old. Kruskal-Wallis test, *P* < 0.0001; from left, *n* = 36, 47, 71. n.s., not significant *P* > 0.05; *, *P* < 0.05; **, *P* < 0.01; ***, *P* < 0.001.

### SF appears in the aged male with an intact gut

To examine the intestinal integrity, aged males were separated into SF and NSF groups, and their gut was stained by anti-Discs large (Dlg) antibody (*12*), which localizes to septate junctions forming gut barrier. Dlg localized to the periphery of individual epithelial cells in both young and aged males with SF, whereas it was reduced in aged males with NSF (Fig. 1K), indicating compromised gut barrier integrity in this group. To further assess the gut barrier integrity, aged males were fed a blue dye during the sleep assay, in which loss of the gut barrier function results in dye-spreading throughout the body (generating ‘Smurf’ flies) (*13*) (Fig. 1L). The gut-intact flies (non-Smurf) showed SF, whereas the leaky-gut flies (Smurf) did not (Fig. 1M), suggesting a negative correlation between gut and sleep homeostasis. The gut integrity influences the gut microbial community (*14–16*), which may be involved in SF. Administration of an antibiotics cocktail from 10-day old, which nearly eradicated the intestinal microbiome (fig. S1K), suppressed SF in aged males (Fig. 1N), demonstrating that microbiome contributes to the development of SF. Of note, aged males with SF exhibited a lower bacterial load compared with aged males with NSF (fig. S1J), suggesting that the bacterial load per se is not a causal factor in SF. These observations led us to hypothesize that variability in the gut homeostasis among aged males permits the selective expansion of certain microbial species, which may predispose a subset of individuals to the SF phenotype.

### *Acinetobacter junii*-derived metabolite induces SF

To characterize microbiome composition in aged males with SF and NSF, we performed 16S rDNA amplicon sequencing for individual flies in each group (Fig. 2A and table 2). Genus-level taxa assignment suggested that *Acinetobacter* could be differentially populated between aged males with SF and NSF (Fig. 2A). The targeted sequence includes sequence variabilities within the same genus, allowing us to narrow down our target species to specific *Acinetobacter* species, including *A. pitti, A. junii, A. baumannii,* and *A. ursingii.* Notably, aged males with SF showed a significant enrichment of *A. junii*, compared with aged males with NSF (Fig. 2A), whereas no such enrichment was observed for other *Acinetobacter* species (fig. S2A). Targeting *A. junii*-specific variable sequence by qPCR further confirmed the increase in the *A. junii* abundance in aged males with SF (Fig. 2B). To test the causal relationship between *A. junii* and SF, the 3-day-old males were fed food mixed with different doses of *Acinetobacter* for 5 days. The young males fed *A. junii,* but not other *Acinetobacter* species, exhibited SF (Fig. 2C). We then tested whether SF is induced by bacterial metabolites or cellular components by feeding either the culture medium or the bacteria debris, respectively. SF was induced by feeding young males the culture media, but not bacterial debris, of *A. junii* (Fig. 2, D and E), suggesting that metabolites produced by *A. junii* are responsible for inducing SF. Feeding the culture media of *A. junii* also induced SF in young females (fig. S2B). Metabolome analyses of the culture media revealed enrichment (fold increase > log_2_ 2.5) of urocanic acid, putrescine, and riboflavin in the media from *A. junii*, compared with those from other *Acinetobacter* species (fig. S2, C to E, and table 3). Among them, urocanic acid induced SF in young males (Fig. 2F). This finding supports the idea that microbiome heterogeneity across individuals, particularly the presence of *A. junii*-derived metabolite is critical for determining age-dependent SF.

**Fig. 2.**
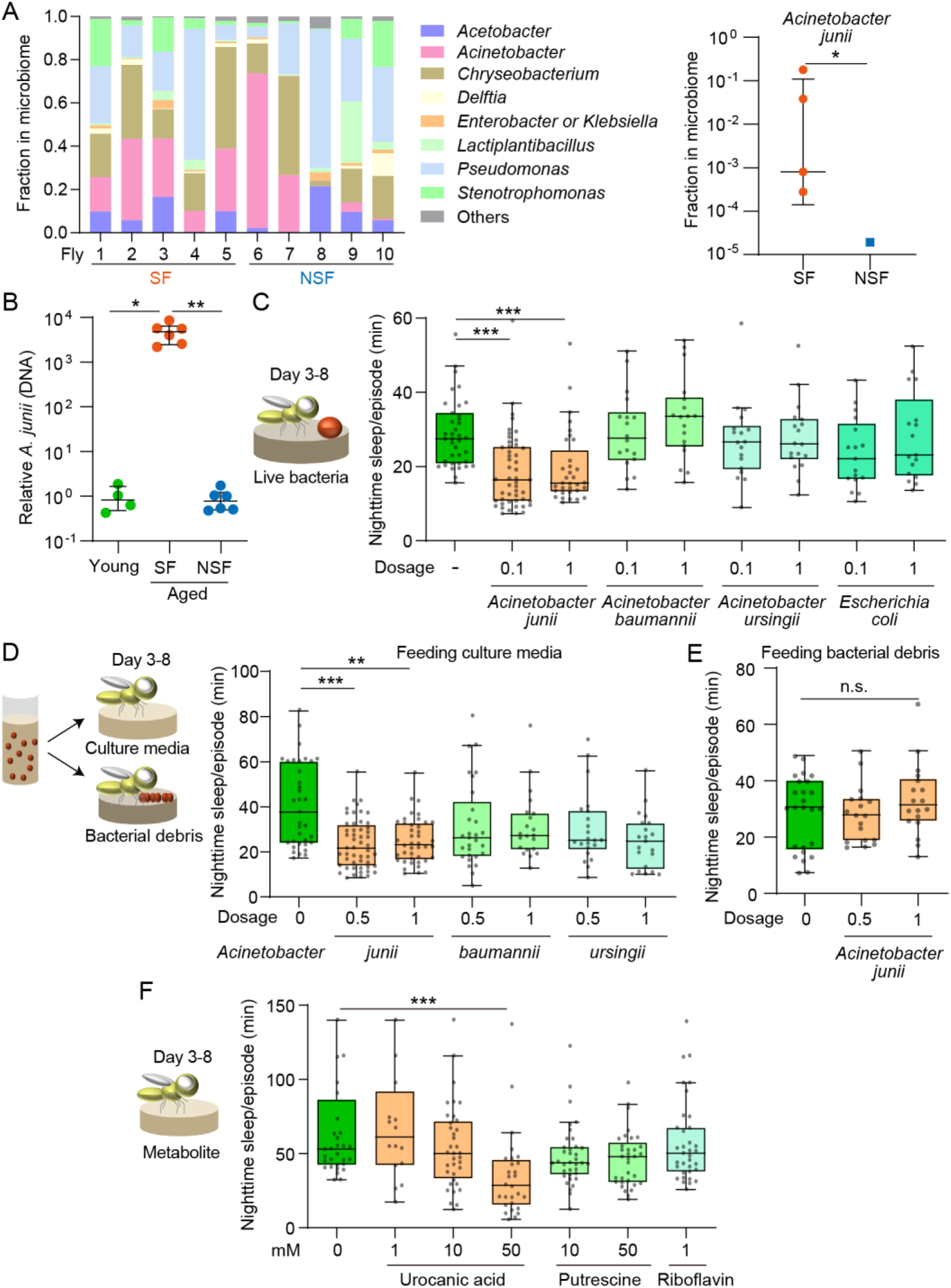
Acinetobacter junii induces SF. **(A)** Microbiome population determined by bacterial 16S rDNA amplicon sequencing. Sleep was analyzed using male flies at 25-day old, and 250 bp of 16S rDNA was amplified by PCR. Genus population determined by sequence of 16S rDNA was shown in five individual flies with SF and NSF. (right) Frequency of read counts determined as *A. junii* was shown. Two-sided Mann-Whitney U-test; *n* = 5 for all, (right) the 5 data points of 0 value were not shown. **(B)** *Acinetobacter junii* is enriched in aged males with SF. DNA was extracted from 10 flies each of the indicated groups, which were analyzed by *A. junii*-specific primers via qPCR. Kruskal-Wallis test *P* < 0.0001; from left, *n* = 4, 6, 6. **(C)** Feeding young flies *Acinetobacter junii* induces SF. Male flies at 3-day old were treated with food containing cultured *Acinetobacter* at the indicated dosages, where dosage 1 indicates 4.6x10^7^ cells/mL. Sleep was analyzed after feeding for 5 days. Kruskal-Wallis test, *P* < 0.0001; from left, *n* = 39, 49, 33, 29, 18, 18, 18, 17, 17. **(D)** Feeding young flies the culture media of *Acinetobacter junii* induces SF. Male flies at 3-day old were treated with liquid food containing the media in which *Acinetobacter* was cultured, at the indicated dosages, where dosage 1 indicates 80 % of the media. Sleep was analyzed after feeding for 5 days. Kruskal-Wallis test, *P* < 0.0001; from left, *n* = 35, 52, 42, 29, 21, 22, 21. **(E)** Feeding young flies dead *Acinetobacter junii* does not induce SF. Male flies at 3-day old were treated with food containing equivalent amount of *Acinetobacter* used in (C) which had been sonicated to exclude the live bacteria. Sleep was analyzed after feeding for 5 days. Kruskal-Wallis test, *P* = 0.6376; from left, *n* = 25, 18, 18. **(F)** Feeding young flies urocanic acid induces SF. Male flies at 3-day old were treated with food containing the indicated metabolites. Sleep was analyzed after feeding for 5 days. Kruskal-Wallis test, *P* < 0.0001; from left, *n* = 32, 16, 36, 28, 36, 33, 35. n.s., not significant *P* > 0.05; *, *P* < 0.05; **, *P* < 0.01; ***, *P* < 0.001.

### Host response mediated by SPH93 from fat body is involved in SF

Our data suggested that *A. junii*-derived urocanic acid is causally linked to SF. However, it remains unclear why flies raised in the same environment could harbor distinct microbiome compositions. This could be attributed to stochasticity in the host’s intestinal environment, competitive dynamics within the microbiota itself, or variability in host–microbe interactions. To explore whether host-microbe interactions contribute to the heterogeneity in *A. junii* abundance, we searched for anti-bacterial response genes that are upregulated in aged males with SF (type A genes). However, transcriptome results from fly bodies did not reveal such genes, particularly for the type A genes (Fig. 1I and table 1). In contrast, a comparison between aged male heads with SF and NSF revealed a greater number of type A genes. Among 20 DEGs exhibiting more than 2-fold difference (Fig. 3, A to C), 9 type A genes were upregulated in SF flies, 7 type B genes were downregulated in NSF flies, 2 type C genes were downregulated in SF flies, and 2 type D genes were upregulated in NSF flies (Fig. 3, D and E, and table 4). Genes of type A are of particular interest, due to the potential causal role in SF. Using available RNAi lines expressing inverted repeats (IR) of the target mRNA, we knocked down each type A gene in the corresponding cell types where the genes are expressed, by referring to single-cell RNA-seq data (*17*). Knockdown of *Serine protease homolog 93* (*SPH93*) via fat body GAL4 (*cg-GAL4*) suppressed SF in aged males (Fig. 3F), whereas knockdown of other genes did not (fig. S3, A to E). We noted that expression of *SPH93* also tends to increase in the aged male bodies with SF compared with those with NSF, although it was not statistically significant (q value = 0.36, 1.44-fold). *SPH93* encodes a serine protease, which may be involved in antibacterial response (*18*). To confirm the expression of *SPH93*, we generated *GAL4*-knock-in flies at the *SPH93* gene locus. The *SPH93::T2A-GAL4* flies carrying *UAS-GFP* showed no obvious GFP expression in young flies, whereas aged flies exhibited significant GFP expression in the fat body marked by Nile red staining (fig. S3F). Importantly, *SPH93* overexpression in the fat body did not induce SF in young flies (fig. S3G), suggesting that *SPH93* is not a direct cause of SF. Although *SPH93*-knockdown suppressed age-dependent SF, it did not affect *A. junii*- or urocanic acid-induced SF in young males (Fig. 3G and S3H), suggesting that SPH93 controls age-dependent changes associated with SF, such as proliferation of *A. junii*. We hypothesized that *A. junii* induces *SPH93* expression to suppress competing bacterial populations and promotes its own expansion, thereby making *SPH93* essential for age-dependent SF. Indeed, feeding young males with *A. junii*, but not other *Acinetobacter* species, induced *SPH93* expression (Fig. 3H). Furthermore, knockdown of *SPH93* increased total bacterial load in aged males (Fig. 3I), while reducing the abundance of *A. junii* (Fig. 3J), supporting a requirement of *SPH93* in *A. junii* propagation. *SPH93*-knockdown increased the proportion of Smurf-positive individuals in aged males (fig. S3I), suggesting that SPH93 also contributed to maintaining gut integrity, most likely by controlling the microbiome. Meanwhile, overexpression of *SPH93* appeared to increase the *A. junii* abundance, although it was not statistically significant (fig. S3J). Together, these results support the notion that *A. junii* exploits host response mediated by *SPH93* expression to facilitate its own proliferation.

**Fig. 3.**
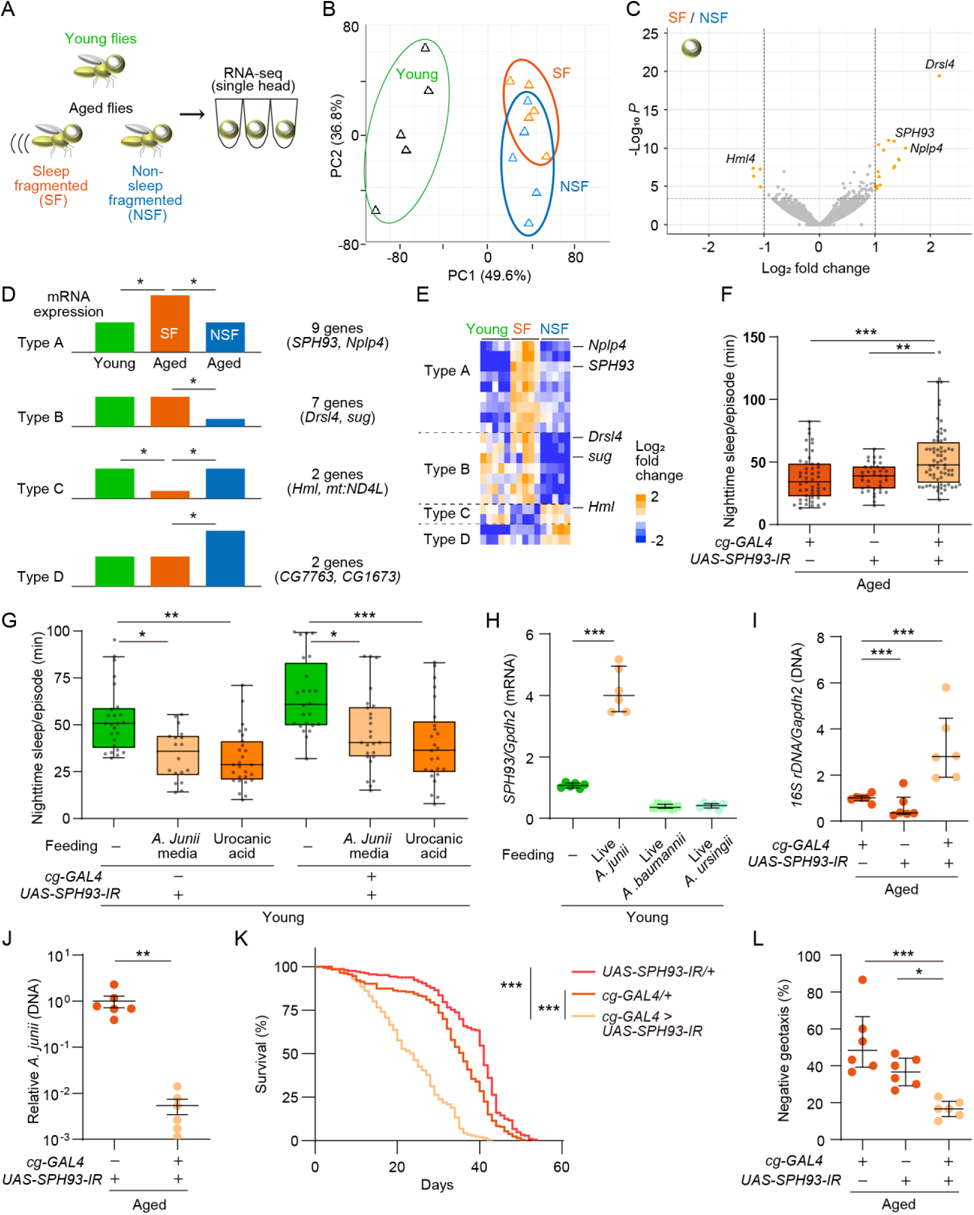
*Acinetobacter junii* exploits host response mediated by *SPH93* to control microbiome. **(A)** Schematics of RNA-seq from individual fly heads. **(B)** Principal component analysis (PCA) of mRNA expression data from the RNA-seq analysis. **(C)** Volcano plot representing the differentially expressed genes (DEG) in fly heads. **(D and E)** Classification (D), heatmap of the log_2_ fold changes in expression (E) of the DEGs. **(F)** Knockdown of *SPH93* suppresses age-dependent SF. The indicated flies at 25-day old were examined by sleep assay. Kruskal-Wallis test, *P* < 0.0001; from left, *n* = 49, 37, 68. **(G)** Knockdown of *SPH93* does not affect SF induced by feeding *A. junii* and urocanic acid. The 3-day-old flies carrying the indicated transgenes were fed food containing 80% of the *A. junii*-cultured medium or 50 mM of urocanic acid for 5 days and examined by sleep assay. Kruskal-Wallis test, *P* < 0.0001; from left, *n* = 25, 24, 18, 25, 27, 25. **(H)** *SPH93* mRNA expression is induced by feeding *A. junii*. The flies at 3-day old were fed food containing cultured live *Acinetobacter*. RNA extracted from the pooled fly heads were analyzed via RT-qPCR. One-way ANOVA, *P* < 0.0001; *n* = 6 for all. **(I)** Knockdown of *SPH93* increases the total bacterial load. The indicated flies at 25-day old were sampled and their bacterial 16S rDNA were analyzed via qPCR. The amount of 16S rDNA was normalized by that of fly genomic *Gapdh2* DNA. One-way ANOVA, *P* < 0.0001; *n* = 6 for all. **(J)** Knockdown of *SPH93* reduces the abundance of *A. junii.* DNA was extracted and pooled from 10 flies each of the indicated groups at 25-day old, which were analyzed by *A. junii*-specific primers via qPCR. Two-sided Mann-Whitney U-test; *n* = 6 for all. **(K and L)** Knockdown of *SPH93* reduces longevity (K) and impairs negative geotaxis at 25-day old (L). (K) Log-rank test, *n* = 200 for all. (L) Kruskal-Wallis test, *P* < 0.0001; *n* = 6 for all. n.s., not significant *P* > 0.05; *, *P* < 0.05; **, *P* < 0.01; ***, *P* < 0.001.

As NSF aged males exhibited shorter lifespan and reduced negative geotaxis, *SPH93*-knockdown male flies exhibited reduced longevity (Fig. 3K) and impaired negative geotaxis after aging (Fig. 3L). Overexpression of *SPH93* extended lifespan (fig. S3K), whereas changes in negative geotaxis were not observed (fig. S3L), indicating a link between SPH93 and other aging phenotypes.

### Upregulation of dopaminergic activity in the mushroom body is required for SF

To further understand the mechanisms underlying age-dependent SF, we conducted targeted metabolomics on aged males with SF and NSF. Out of 144 water-soluble metabolites including neurotransmitters, only dopamine levels were significantly elevated in aged males with SF, showing an increase of > 1.5-fold (Fig. 4A and table 5). Because dopamine upregulation promotes wakefulness (*19, 20*), we tested whether dopamine signaling is involved in age-dependent SF using dopamine receptor mutants. Dopamine type 1 receptors are encoded by *Dop1R1* and *Dop1R2*, while the dopamine type 2 receptor is encoded by *DopR2*. The *Dop1R2* and *DopR2* mutants, but not *Dop1R1* mutants, significantly suppressed SF in aged males (Fig. 4B). Similarly, pan-neuronal knockdown of *Dop1R2* and *DopR2* using *nSyb-GAL4* suppressed SF in aged males (fig. S4, A and B). Dopamine neurons innervate several neuropiles (*21*), which have been associated with sleep regulation, including the ellipsoid body (EB) (*22*), the fan-shaped body (FSB) (*23, 24*), and the mushroom body (MB) (*25, 26*). SF in aged males was suppressed by the knockdown of *Dop1R2* and *DopR2* in the MBs (Fig. 4, C and D), but not in the EB or FSB (fig. S4, C to F). Dopaminergic activity can be assessed using DopR-TANGO flies (*27*) in which dopamine binding to the artificial TANGO receptor induces expression of a reporter gene, myr-tdTomato. The reporter expression in the MB lobes was higher in aged males with SF than in those with NSF (Fig. 4E), indicating upregulation of dopaminergic activity. Taken together, upregulation of dopaminergic activity in the MBs plays a crucial role in inducing age-dependent SF.

**Fig. 4.**
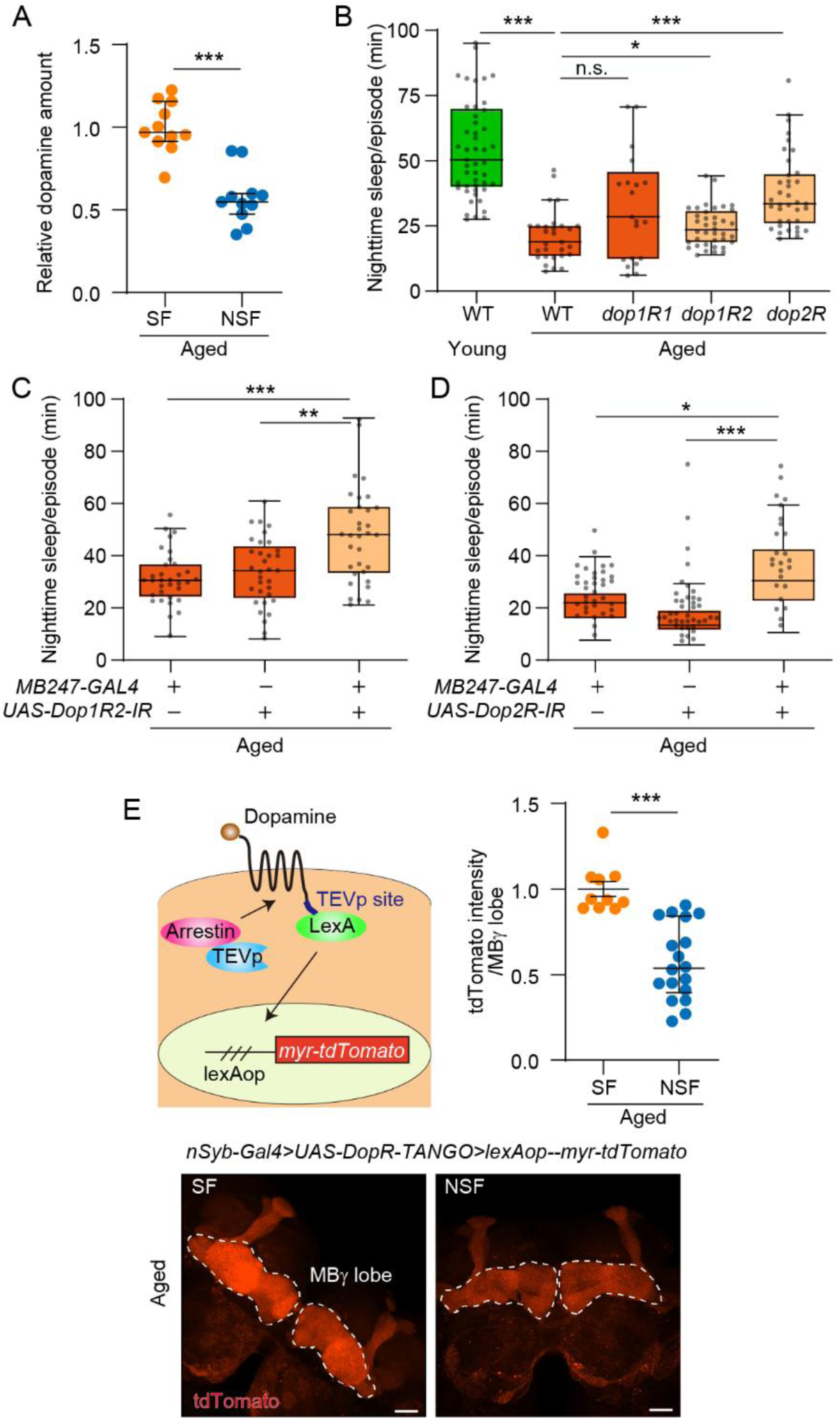
Upregulation of dopaminergic activity is required for SF. **(A)** Dopamine amount is increased in aged males with SF. Sleep was analyzed using aged males at 25-day old, and the heads from flies showing SF or NSF were pooled. The head extracts were analyzed by LC-MS/MS. Two-sided Mann-Whitney U-test; *P* < 0.001 *n* = 11 for all. **(B)** *Dop2R* and *Dop1R2* are involved in age-dependent SF. Sleep was analyzed using wild-type (WT) flies, mutant flies in *Dop1R1* (*dumb^2^*), *Dop2R* (*dop2R-KO*), and *Dop1R2* (*damb*) at 5-day or 25-day old. Kruskal-Wallis test, *P* < 0.0001; from left, *n* = 49, 29, 21, 37, 38. **(C and D)** *Dop1R2* (C) and *Dop2R* (D) in MB neurons are involved in age-dependent SF. Sleep was analyzed using the indicated flies at 25-day old. (C) Kruskal-Wallis test, *P* < 0.0001; from left, *n* =33, 33, 33. (D) Kruskal-Wallis test, *P* < 0.0001; from left, *n* = 35, 40, 25. **(E)** Dopaminergic activity is increased in aged males with SF. Sleep was analyzed using the indicated flies, and the flies with SF or NSF were dissected and fixed. The fluorescence raw signal of tdTomato was quantified from MB γ neurons (white dotted line). Scale bar: 20 μm. Two-sided Mann-Whitney U-test; *n* = 10, 18. n.s., not significant *P* > 0.05; *, *P* < 0.05; **, *P* < 0.01; ***, *P* < 0.001.

### Urocanic acid increases dopaminergic activity through crosstalk with a serotonin receptor

The significance of dopamine upregulation in SF prompted us to test whether *A. junii* and urocanic acid have causal roles in increasing dopaminergic activity. Feeding young males the culture media of *A. junii* or urocanic acid elevated the DopR-TANGO reporter expression (Fig. 5A), demonstrating that *A. junii*-derived urocanic acid indeed enhances dopaminergic signaling. Furthermore, SF in young males, induced by feeding the culture media of *A. junii* or urocanic acid, was suppressed in *dop1R2* and *dopR2* mutant flies (Fig. 5, B and C), although the suppression of SF caused by *A. junii* media in *dopR2* mutant flies was not statistically significant (Fig. S5, A and B). These results imply that urocanic acid directly or indirectly augments dopaminergic activity to induce SF. The precursor of urocanic acid is histidine, which is also a precursor of histamine known to maintain wakefulness (*28*). Urocanic acid and histamine are structurally similar, potentially leading to activation of the histamine receptor. However, SF in aged males was not affected by a histamine receptor antagonist, hydroxyzine (Fig. 5D). Urocanic acid also acts as an agonist of a mammalian serotonin receptor, 5-HT2A (*29*), raising the possibility that urocanic acid induces the activation of serotonin receptors to upregulate dopaminergic activity, thereby causing SF. Indeed, feeding a serotonin receptor antagonist, methysergide maleate, suppressed SF in aged males (Fig. 5E). There are five genes that encode serotonin receptors in flies. Pan-neuronal knockdown of a serotonin receptor encoded by *5-HT2A* using *nSyb-GAL4*, but not the other receptors, suppressed SF in aged males (Fig. 5F, fig. S5, C to F). Knockdown of *5-HT2A* also suppressed the reporter expression of DopR-TANGO induced by aging (Fig. 5G), feeding the culture media of *A. junii* (fig. S5G), and feeding urocanic acid (fig. S5H), suggesting that upregulation of dopaminergic activity by urocanic acid requires 5-HT2A. Furthermore, knockdown of *5-HT2A* by dopamine neuron *GAL4* suppressed SF in aged flies with a pronounced effect of *Ddc-GAL4* labeling the protocerebral anterior medial (PAM) dopamine neurons, rather than *TH-GAL4* labeling the protocerebral posterior lateral (PPL) dopamine neurons (*30*) (Fig. 5H). Consistently, SF in young flies induced by feeding *A. junii* or urocanic acid was suppressed by knockdown of *5-HT2A* via *Ddc-GAL4* (fig. S5, I and J). By employing transgenic flies carrying the C-terminus of split GFP (spGFP_11_) and an HA tag at the endogenous *5-HT2A* gene (*31*), particularly at the stop codons of two isoforms (*5-HT2A-S* and *-L*), we tested the expression of 5-HT2A in dopamine neurons. The N-terminus of spGFP (spGFP_1-10_) was expressed using *Ddc-GAL4*. Although 5-HT2A-L proteins were not detectable, 5-HT2A-S proteins were detected in *Ddc-GAL4*-positive neurons (Fig. 5I). We noticed that anti-HA antibody staining produced detectable signals rather than the reconstituted GFP. Taken together, urocanic acid-dependent upregulation of dopaminergic activity to induce SF is mediated by a serotonin receptor, 5-HT2A.

**Fig. 5.**
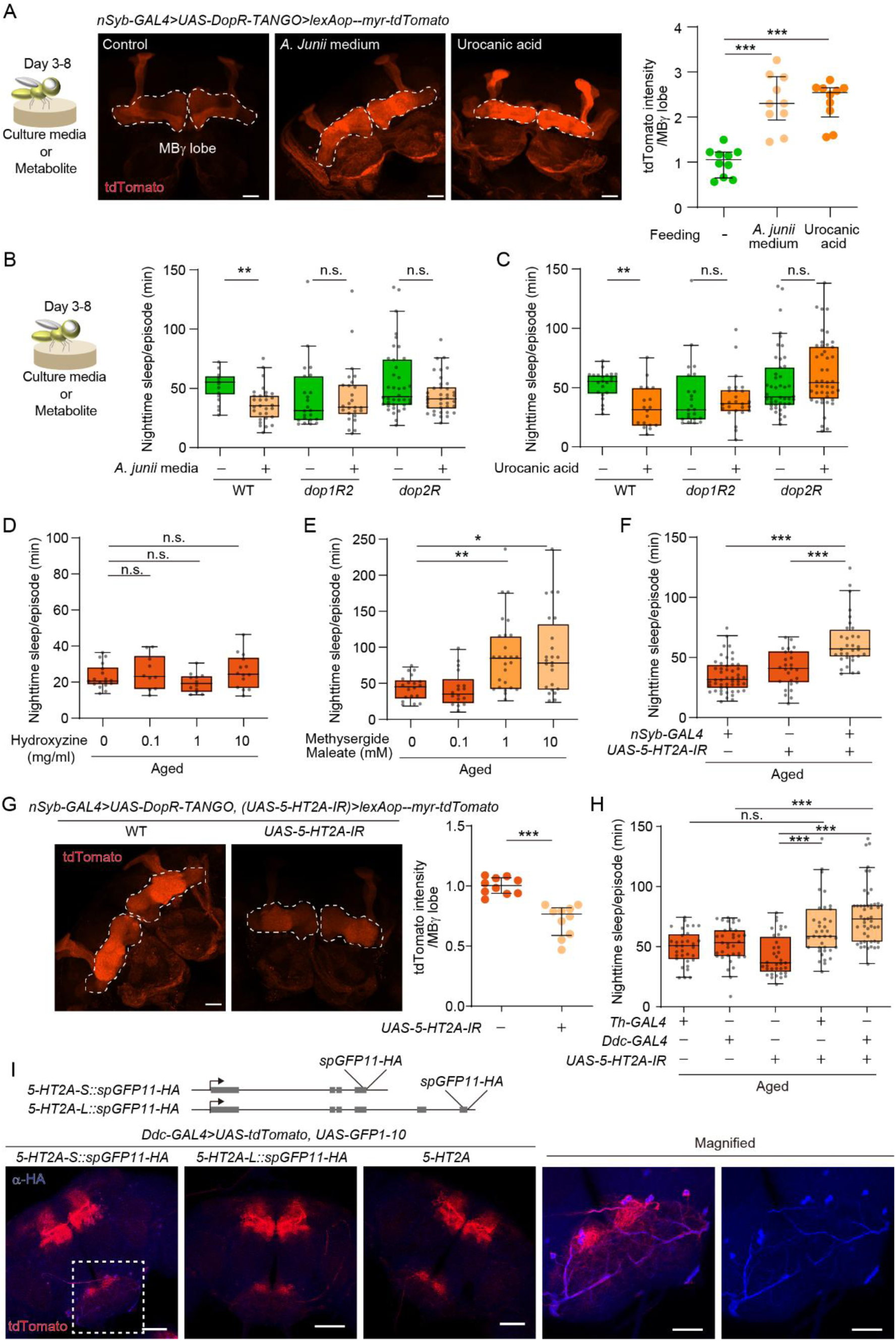
*A. junii* dependent upregulation of dopamine is mediated by serotonin receptor 5-HT2A. **(A)** Feeding *A. junii* or urocanic acid increases dopaminergic activity in young males. The 3-day-old flies with the indicated transgenes were fed liquid food containing 80% of the *A. junii*-cultured medium or 50 mM of urocanic acid for 5 days. The dissected brains were fixed and the fluorescence raw signal of tdTomato was quantified from MB γ neurons (white dotted line). Scale bar: 20 μm. Kruskal-Wallis test, *P* < 0.0001; *n* = 10 for all. **(B and C)** *Dop1R2* and *DopR2* are required for SF induced by feeding *A. junii* media or urocanic acid. Wild-type (WT) flies and the mutant flies in *Dop1R2* (*dop1R2*) or *Dop2R* (*dop2R*) at 3-day old were fed liquid food containing 80% of the *A. junii*-cultured medium (B) or 50 mM of urocanic acid (C) for 5 days and examined by sleep assay. (B) Kruskal-Wallis test, *P* < 0.0001; from left, *n* = 22, 30, 23, 26, 38, 38. (C) Kruskal-Wallis test, *P* < 0.0001; from left, *n* = 22, 18, 23, 26, 45, 45. **(D and E)** Age-dependent SF is not suppressed by a histamine receptor antagonist (D) but suppressed by a 5-HT receptor antagonist (E). Aged males at 22-day old were treated with food containing the indicated doses of histamine receptor antagonist, hydroxyzine (D) or serotonin receptor antagonist, methysergide maleate (E) for 3 days, and examined by sleep assay. (D) Kruskal-Wallis test, *P* = 0.2807; from left, *n* =18, 10, 12, 14. (E) Kruskal-Wallis test, *P* < 0.0001; from left, *n* = 19, 17, 26, 24. **(F)** Knockdown of *5-HT2A* suppresses age-dependent SF. The indicated flies at 25-day old were examined by sleep assay. Kruskal-Wallis test, *P* < 0.0001; from left, *n* = 50, 26, 32. **(G)** *5-HT2A* is required for upregulation of dopaminergic activity in aged flies. The 25-day-old flies with the indicated transgenes were dissected and the brains were fixed. The fluorescence raw signal of tdTomato was quantified from MB γ neurons (white dotted line). Two-sided Mann-Whitney U-test; *n* = 10 for all. **(H)** 5-HT2A expression in dopamine neurons is required for SF. Flies at 25-day old were examined by sleep assay. Kruskal-Wallis test, *P* < 0.0001; from left, *n* = 25, 34, 27, 27, 34. **(I)** Short isoform of *5-HT2A* is expressed in dopamine neurons. Flies with indicated transgene were dissected, brains were fixed, and the fluorescence signals were detected using anti-tdTomato (red) and anti-HA antibody (blue) in SEZ area (white dotted line). Scale bar: (left) 50 μm, (right) 20 μm. n.s., not significant *P* > 0.05; *, *P* < 0.05; **, *P* < 0.01; ***, *P* < 0.001.

### Urocanic acid-producing bacteria extend lifespan in a SPH93-depnedent manner

Given that aged males with SF live longer, we hypothesized that urocanic acid-producing bacteria also affect lifespan. Feeding flies *A. junii*, but not *A. baumannii*, starting at 5-day old, slightly but significantly extended their lifespan (Fig. 6A). To examine the significance of urocanic acid, the gene encoding its biosynthetic enzyme, histidase (*hutH*) (*32*), was cloned from *A. junii* into *Escherichia coli*, which was fed to the flies. This resulted in extended lifespan (Fig. 6B), and induced SF in young flies (Fig. 6C). In contrast, feeding the culture media of *A. junii* did not induce lifespan extension (Fig. 6D), even though it induced SF (Fig. 2D), suggesting that lifespan extension requires the bacteria producing urocanic acid, rather than urocanic acid alone. Knockdown of *SPH93* in the fat body suppressed the lifespan extension induced by feeding *A. junii* or *hutH*-expressing *E. coli* (Fig. 6, E and F), suggesting that *SPH93* is required to mediate the beneficial effect on lifespan. Thus, urocanic acid-producing bacteria induce lifespan extension by modulating host gene expression, *SPH93*.

**Fig. 6.**
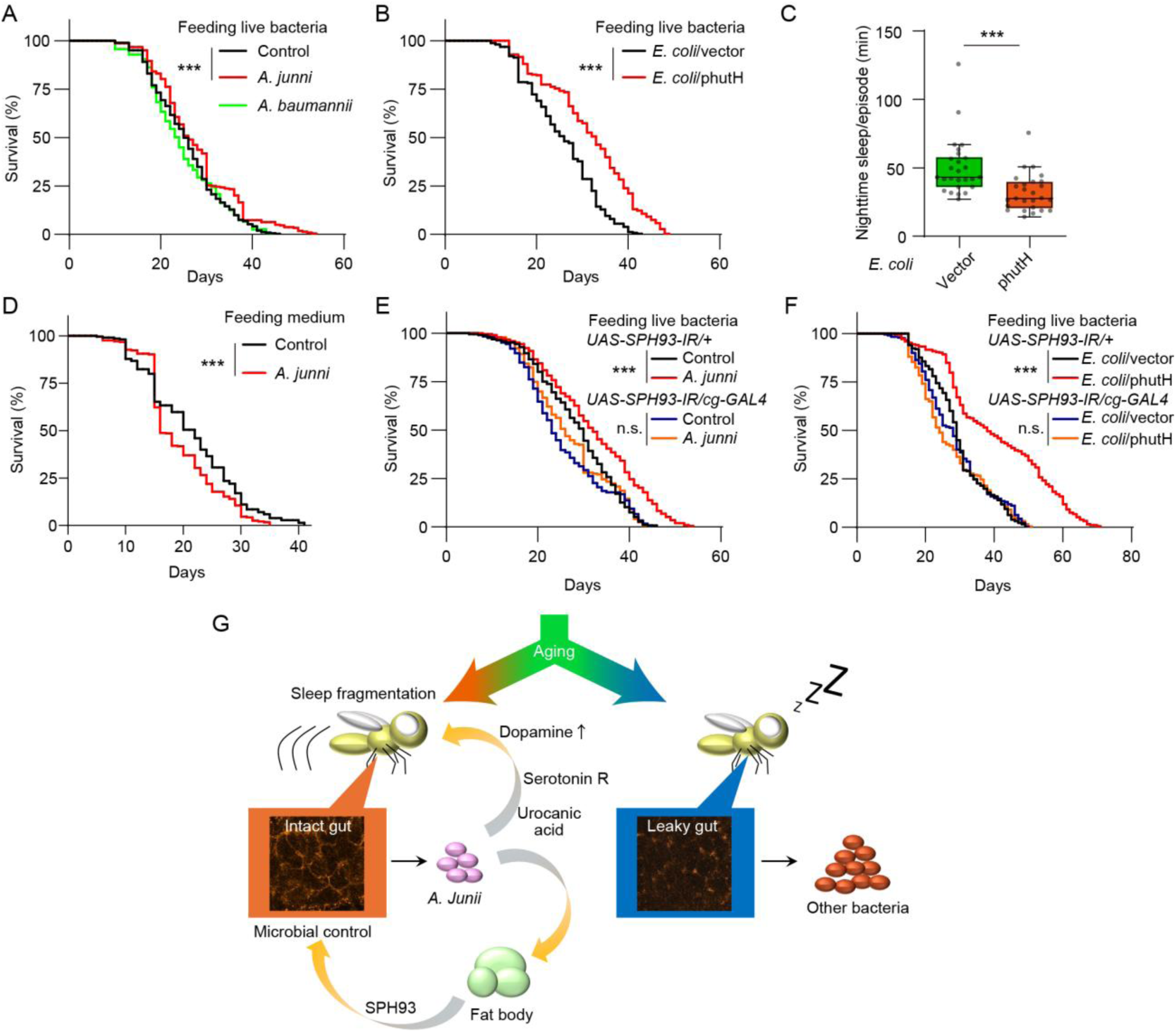
Urocanic acid-producing bacteria extend lifespan dependent on SPH93. **(A and B)** Feeding live *A. junii* (A) or *hutH*-expressing *E. coli* (B) extends lifespan. Wild-type flies were fed the indicated bacteria strains, starting at 5-day old. Log-rank test, (A) *n* = 300 for all; (B) *n* = 300 for all **(C)** Feeding *hutH*-expressing *E. coli* induces SF in young flies. The indicated bacteria were fed flies at 5-day old, and sleep was analyzed at 8-day old. Two-sided Mann-Whitney U-test; *n* = 36, 35. **(D)** Feeding *A. junii*-cultured medium does not affect lifespan. Log-rank test, *n* = 300 for all. **(E and F)** Lifespan extension by feeding *A. junii* (E) or *hutH*-expressing *E coli* (F) requires *SPH93* expression. Log-rank test, *n* =300 for all. **(G)** Model generating heterogeneity in age-dependent SF. Aged flies exhibit variability in gut environment regarding gut homeostasis and microbiome. Propagation of *A. junii* results in circulation of urocanic acid, cross-reacting with a serotonin receptor, 5-HT2A, and the subsequent upregulation of dopaminergic activity, leading to SF. *A. junii* also induces *SPH93* expression in fat body, controlling microbiome, and positively affect lifespan. n.s., not significant *P* > 0.05; *, *P* < 0.05; **, *P* < 0.01; ***, *P* < 0.001.

## Discussion

Aging exhibits significant heterogeneity, influenced by fluctuations in biological processes, individual experiences such as eating habits, and external factors. A comprehensive understanding of individual aging requires a more nuanced analysis of phenotypic variability among the aged population, rather than relying solely on comparisons between young and aged animals. By separating aged individuals based on sleep phenotype, this study revealed that a specific microbe, *A. junii*, is causally related to age-dependent SF. We further provided a neuronal mechanism linking *A. junii* to SF, mediated by the *A. junii* metabolite, urocanic acid, and the upregulation of dopamine signaling. Importantly, in parallel with SF, *A. junii* positively impacts aging through the activation of the *SPH93* pathway of the host, demonstrating a trade-off between lifespan and sleep quality. The expression of *SPH93* is required for populating *A. junii*, indicating that *A. junii* employs the host response for its own proliferation. Thus, our approach to segregating the aging trait revealed a coordinated, system-wide regulation of aging, that integrate the gut, microbiome, host response, and nervous system.

### Aging heterogeneity via microbiome

Dysbiosis associated with aging is generally considered to be accompanied by a decrease in beneficial bacteria (*33*). Flies also exhibit age-related dysbiosis (*34*), and both positive (*34, 35*) and negative (*36*) effects of commensal bacteria on longevity have been reported. Despite the relatively solid concept of dysbiosis related to aging, how individual microbiome traits shape multiple aging phenotypes is still unknown. Our study demonstrated that aged flies host distinct microbiomes with or without *A. junii*, leading to aged populations exhibiting different phenotypes: one population is long-lived with SF and intact gut, and the other is short-lived without SF but with leaky gut. A metabolite from *A. junii*, urocanic acid, plays a central role in this trade-off through direct and indirect routes. SF caused by urocanic acid through 5-HT2A would be direct, as its mammalian homologous receptor 5-HT2A is directly activated by urocanic acid (*29*). Indirectly, urocanic acid extends lifespan, as feeding the cultured medium of *A. junii* alone did not alter longevity (Fig. 6D) while feeding urocanic acid-producing bacteria did promote lifespan extension (Fig. 6, A and B). Urocanic acid is an intermediate metabolite in the synthesis of glutamate from histidine (*32*). We speculate that urocanic acid-producing bacteria utilize urocanic acid or other intermediate metabolites, such as 4-imidazolone-5-propionate and N-formimino glutamate (*32*), to generate the components which induces SPH93 expression, ultimately leading to lifespan extension. Given that SPH93 regulates the overall bacterial load (Fig. 3I), the longevity benefits conferred by urocanic acid-producing bacteria could arise from modulating the abundance or composition of the microbiota through *SPH93* expression. It will be important to characterize the signaling pathway of *SPH93* expression to understand its working mechanism and explore its potential for aging intervention.

SPH93 belongs to the S1A serine protease family (*18*), which is commonly activated by proteolytic cleavage and plays various roles ranging from the digestion of dietary proteins to blood coagulation (*37*). However, SPH93 lacks the His-Asp-Ser catalytic triad, which is a conserved enzymatic domain (*18*), and is therefore named a serine protease homolog. Serine protease homolog proteins are predicted to act as competitors of active proteases or to perform additional functions (*38*). Different serine protease homologs have been reported to be involved in axon guidance (*39*), egg-laying behavior (*40*), and Toll signaling activation (*41*) in flies, and in the migration of sperm in mice (*42*), although detailed molecular mechanisms remain unexplored. Given that SPH93 is required for populating *A. junii* (Fig. 3J) and suppressing the bacterial load in the fly gut microbiota (Fig. 3I), it will be essential to study how SPH93 shapes the microbiome community and how specific microbiome components can adapt to the SPH93-regulated microenvironment.

### Potential mechanisms generating aging heterogeneity

Although *A. junii* is involved in aging heterogeneity in our study, the core mechanisms generating the heterogeneity has yet to be determined. We speculate that the heterogeneity could be attributed to either the microbiome community or the host immune response. Firstly, metabolic co-dependence and competition among bacterial species could generate fluctuations in the microbiome composition (*43*). Secondly, the host response including *SPH93* expression could be variable, therefore diversifying the microbiome composition in individual flies (*44*). Alternatively, rather than considering microbial communities to be stableover time, their heterogeneity may be more dynamic, fluctuating with the type of food consumed and the timing of digestion, such as whether nutrient intake occurs during wakefulness or near the sleep period.

Given that serotonin is involved in sleep regulation (*45*), and the serotonin receptor 5-HT2A in mammals cross-reacts with urocanic acid, urocanic acid-producing bacteria could also affect sleep patterns in mammals. *A. junii* is also reported in human fecal specimens (*46*), although other urocanic acid-producing bacteria may be involved. Recent study on anti-PD-1 therapy for cancer treatment demonstrated that urocanic acid from the microbe, *Muribaculum* improves the efficacy of anti-PD-1 treatment (*47*). In parallel with our observations, *E. coli* expressing *hutH* also recapitulated the beneficial effect (*47*), suggesting broader potential of leveraging the urocanic acid-producing bacteria in mammalian systems.

Overall, our study demonstrated an unanticipated trade-off between lifespan and sleep quality during aging. Given that reduced quality of sleep negatively impacts physical, cognitive, and mental functioning, this trade-off was unexpected. However, age-dependent deterioration in organismal functions could be variable across molecular pathways, cell types, and organs; such trade-off could widely exist yet remains to be characterized. The short lifespan in *Drosophila* allowed us to capture a simplified and robust interaction between lifespan and sleep. Further understanding of aging heterogeneity, including phenotypic trade-offs as such, would require targeted analysis involving different phenotypes and biomarkers. Such efforts will inform strategies to mitigate trade-offs, ultimately promoting healthy aging.

## Materials and Methods

### Fly and bacterial strains

Our wild-type control line, *w(CS10)* (*48*), was used for most behavioral experiments or as a control unless otherwise stated. The *Dop1R1* (*dumb2*) (*49*) and *Dop1R2* (*damb*) mutant fly lines (*50*) and *TH-GAL4* (*51*) flies were obtained from M. Saitoe (Tokyo Metropolitan Institute of Medical Science, Japan). *DopR2-KO* flies were obtained from Yi Rao (*52*) (Peking University, China). *Cg-GAL4* was obtained from Y. Yan (The Hong Kong University of Science and Technology, Hong Kong SAR, China). Other *GAL4* lines including *nSyb-GAL4* (39171), *Ddc-GAL4* (7010), *FSB-GAL4* (*R75G12-GAL4*, 39906), *MB247* (50742), *EB-GAL4* (*R15F02-GAL4*, 48698) and *5-HT2A-S::spGFP11-HA* (605062), *5-HT2A-L::spGFP11-HA* (605061), and *UAS-DopR-tango* (68234) were obtained from Bloomington Drosophila Stock Center (Indiana, USA). *UAS-SPH93-IR* (KK104307), *UAS-Dop1R1-IR* (KK102341), *UAS-Dop1R2-IR* (KK105324), *UAS-Dop2R-IR* (GD11471), *UAS-Nplp4-IR* (KK104662), *UAS-CG9312-IR* (GD16646), *UAS-CG43092-IR* (KK102964), *UAS-CG34227-IR* (GD12520), *UAS-CG13067-IR* (KK103334), *UAS-5-HT1A-IR* (KK106094), *UAS-5-HT1B-IR* (KK109929), *UAS-5-HT2A-IR* (KK102105), *UAS-5-HT2B-IR* (KK102356), and *UAS-5-HT7-IR* (KK104804) were obtained from Vienna *Drosophila* RNAi Center (Vienna, Austria). All fly strains were outcrossed at least 5 to 10 times with our wild-type strains, *w(CS10)*. The *SPH93::T2A-GAL4* knock-in and *UAS-SPH93* transgenic flies were obtained via germline transformation using standard procedures. Bacterial strains *Acinetobacter junii* (110497)*, Acinetobacter baumannii* (110490), and *Acinetobacter ursingii* (110511) were obtained from NRRC (Japan).

### Plasmid construction

To obtain the *SPH93::T2A-GAL4* flies via CRISPR/Cas9, two plasmids were constructed. Firstly, the pCFD4-SPH93 expressing two sgRNA targeting *SPH93* nearby the stop codon was obtained by amplifying a fragment containing two sgRNA via PCR and cloning it into pCFD4 (Addgene no. 49411), as previously described (*53*). To construct the donor plasmid, pSPH93::T2A-GAL4, T2A sequence was cloned into the XhoI-BglII-digested pBluescript SK-(-) vector, resulting in pT2A. GAL4.1 coding region from pBPGAL4.1Uw (Addgene no. 26226) was cloned into the BglII-XbaI-digested pT2A, resulting in pT2A-GAL4.1. DNA fragments at the length of 0.7 kb franking the stop codon of *SPH93* were amplified by PCR, and cloned into KpnI/XhoI and NotI/SacI sites in pT2A-GAL4.1, resulting in pSPH93::T2A-GAL4.

To construct UAS-SPH93, the coding region of *SPH93* was amplified from cDNA by PCR, and cloned into the NotI-XbaI-digested pJFRC-MUH (Addgene no. 26213). The *hutH* over-expression plasmid, phutH, was constructed using Gibson assembly (*54*). A tac-promoter containing plasmid (pLW212) backbone was linearized by PCR amplification using primers designed to retain the tac promoter and plasmid origin. The *hutH* gene was amplified from the *Acinetobacter junii* (NBRC strain 110497) genome, and the kanamycin resistance (KanR) cassette was amplified from pUA66-theta-GFP (*55*). The obtained PCR fragments were assembled using NEBuilder HiFi DNA Assembly Master Mix (New England Biolabs Inc., Ipswich, MA, USA), resulting in phutH.

To construct UAS-spGFP_1-10_, the GFP-encoding fragment was amplified by PCR, lacking the C-terminal 16 amino acids, and cloned into the BglII-XbaI-digested pUAST plasmid (*56*).

### Fly culture conditions

Flies were raised under a 12-h light:dark cycle, at a temperature of 25 °C and humidity of 55±5%. Adult flies were collected under carbon dioxide anesthesia in groups of 50-70 flies in food vials, and for behavioral experiments, males and females were separated under carbon dioxide anesthesia at least 2 days before test. Hydroxyzine dihydrochloride (H1832, TCI, Tokyo, Japan), Methysergide maleate (5.06419. Sigma, St. Louis, MO, USA), Riboflavin 5’-phosphate sodium salt hydrate (R7774, Sigma, St. Louis, MO, USA) and Putrescine dihydrochloride (29428-01, nacalai tesque, Kyoto, Japan) were mixed in 2.5% sucrose and 2.5 % bacto yeast extract (212750, Thermo Fisher Scientific, San Jose, CA, USA) and administered to the flies on Whatman 3MM filter paper for 3-5 days. Urocanic acid (36008-81, nacalai tesque, Kyoto, Japan) was dissolved in 0.2 M NaOH at the final concentration of 1-50 mM, and mixed in 2.5% sucrose and 2.5 % bacto yeast extract and administered to the flies on Whatman 3MM filter paper for 3-5 days. As a control experiment, 0.2 M NaOH was added to the liquid food. The antibiotic-fed flies were maintained on autoclaved solid food supplemented with an antibiotic cocktail containing ampicillin (BP17605, Fisher Bioreagents, MA, USA), tetracycline (T3258, Sigma, St. Louis, MO, USA), and kanamycin at a concentration of 50 µg/mL.

### Bacterial feeding

To feed flies *E. coli* carrying phutH, *E. coli* was cultured overnight in LB medium supplemented with 50 µg/mL kanamycin (K1377, Sigma, St. Louis, MO, USA) at 37°C. The culture was diluted at 1:100 in a fresh LB medium containing 50 µg/mL kanamycin and incubated at 37°C until reaching to a density of OD₆₀₀ 0.5–0.6. Isopropyl β-D-1-thiogalactopyranoside (IPTG) (GEN-S-02122-25G, ProteinArk, Sheffield, UK) was added to a final concentration of 1 mM, and incubated for more 10-12 hours at 18°C. The bacteria obtained were spotted on the fly solid food containing 1 mM IPTG. Other bacteria, including *A. junii*, were cultured in LB medium at 37°C and fed flies on the normal solid food. Bacterial culture media was dissolved in 2.5 % sucrose, which was soaked on Whatman filter paper prior to feeding the flies. Cultured bacteria were sonicated to feed dead bacteria.

### Sleep assay

The arena to monitor sleep was 3D-printed using white PLA material (fig. 1A) with 20 mm diameter and 7 mm height. Each well contains a reservoir, filled with 10 mL of 2.5% sucrose and 2.5 % bacto yeast extract, and the floor covered with 1% agarose. The individual flies were gently loaded using an orally operated fly aspirator into each circular arena at ZT10, and recording for 36 hours was started at ZT12 under a 12-h light:dark cycle. Analyses excluded the first evening peak and performed after ZT13 and onward. Behavioral recordings were conducted inside a custom-made acrylic box at a temperature of 25°C and humidity of 55 ± 5%. A Basler camera (acA1300-60gmNIR GigE), paired with a Kowa lens (LM12HC-SW), was mounted above the arena. White light was placed inside the box under a 12-h light:dark cycle, while infrared (IR) light was illuminated as a backlight to track fly locomotion. An IR long-pass filter (MidOpt LP815-35.5) was placed on the camera lens to block the visible light. The recordings were captured at 15 frames per second using Noldus EthoVision-XT15 or MediaRecorder4.0 image acquisition software. The tracking of flies and behavior analyses was performed by EthoVision XT15 according to the manufacturer’s instruction (Noldus Information Technology). Individual flies showing more than 90% successful tracking were analyzed. The flies which died during the recording was excluded from the analyses. Movement in the tracking system was defined as the fly moving ≥2.0 mm/s, while termination of movement was defined as the fly’s activity dropping below ≤1.5 mm/s. Sleep was calculated by measuring inactivity for longer than 5 min. Ethovision tracking data has been used to calculate total sleep, sleep episodes and average sleep time/episode during daytime (ZT1-11) or nighttime (ZT13-23). The reduction in sleep duration after feeding *A. junii* media or urocanic acid (Dsleep time/episode) was calculated by subtracting the average sleep duration of the control feeding group from each individual data point.

### Smurf assay

Smurf assay was performed as previously described with some modifications (*13*). Flies were fed liquid food containing 2.5% sucrose and 2.5 % bacto yeast extract with Brilliant blue FCF (027-12842, Fujifilm, Wako Pure Chemical Corporation, Japan) at a final concentration of 2.5% for overnight. Flies exhibited blue color in their abdomens were counted.

### Climbing assay

Climbing behavior was analyzed as previously described (*56*) with slight modifications, which was conducted at 25 °C and 55 ± 5% humidity, under permanent light from the top. A group of age-matched 20-25 male flies were loaded to a plastic vial with a hight of 20 cm. The plastic vials were placed in a metal rack, which was sharply dropped to locate the flies at the bottom. After 20 sec, the flies climbing over 15 cm were counted. This was repeated 3 times with 30 sec intervals, and average of three experiments was plotted.

### Longevity assay

The longevity assay was carried out as previously described (*57*). Fifty male flies at 5-day old were collected into vials containing the standard fly food or ones supplemented with live bacteria or media. Flies were transferred to a fresh food vial every 2 days, while the number of dead flies was counted daily. The assay was continued until all flies were dead. Replicates of 4-6 vials were summed to draw the survival curve.

### Immunohistochemistry

Staining was carried out as previously described (*58*). Flies were briefly washed with ice-cold ethanol, which was then replaced with ice-cold phosphate buffered saline (PBS). To collect intestines, the fly’s genital region was pulled gently to remove the digestive system from rest of the fly. For head, the proboscis was removed in PBS, following which the fly heads and intestines were fixed in 4% paraformaldehyde in PBS for 20 min at room temperature. The dissected brains and intestines were treated with PBS containing 0.3% Triton X-100 for 3 h at room temperature and blocked for 1 h in PBS containing 0.1% Triton X-100 and 4% Block Ace (Sumitomo Dainippon Pharma, Osaka, Japan). The samples were then incubated with primary antibodies in blocking solution for overnight at 4°C. After washing three times with PBS containing 0.1% Triton X-100, the brains were incubated with secondary antibodies in blocking solution for 4 h at room temperature. After washing three times with PBS containing 0.1% Triton X-100, the brains were mounted in PermaFluor (Lab Vision Corp., Fremont, CA, USA), and images were captured using a confocal microscope, LSM980 (Zeiss Microsystems, Jena, Germany). The following primary antibodies were used: mouse monoclonal mouse anti-dlg (4F3, DSHB, 1:25), rat anti-tdTomato (EST203, Kerafast, 1:100), mouse anti-HA antibody (26183, Thermo Fisher Scientific, San Jose, CA, USA, 1:1000) and chicken anti-GFP antibody (ab13970, Abcam, Cambridge, MA, USA, 1:1000). Nile red has been purchased from (N3013, Sigma, St. Louis, MO, USA) and used at a concentration of 0.5 µg/mL. The following secondary antibodies were used: donkey anti-chicken immunoglobulin Y (IgY) Alexa Fluor 488, (703-545- 155, Jackson ImmunoResearch Labs, Inc., West Grove, PA, USA), anti-mouse IgG Alexa Fluor 488 (715-545-150, Jackson ImmunoResearch Labs), anti-mouse IgG Alexa Fluor 555 (A-21424, Invitrogen) and anti-Rat IgG Alexa Fluor 647 (A-21236, Invitrogen). Secondary antibodies were diluted to 1:500 in blocking solution, together with DAPI at 2.5 ug/ml.

### RNA sequencing library preparation

Individual heads were collected and homogenized using TRIzol reagent (Thermo Fisher Scientific, San Jose, CA, USA). RNA was extracted with chloroform, ethanol-precipitated, and dissolved in 6 µL diethylpyrocarbonate (DEPC) water. DNA was digested with DNase, and mRNA was further collected using Oligo d(T)25 Magnetic Beads (S1419, New England Biolabs inc., Ipswic(H) MA, USA), according to the manufacturer’s instructions. The obtained mRNA was then used to generate a library with the NEBNext Ultra II RNA Library Prep Kit for Illumina (New England Biolabs Inc., Ipswich, MA, USA), following the manufacturer’s instructions. Paired-end reads were generated on a HiSeq X Ten (Illumina, San Diego, CA, USA).

### 16S rDNA sequencing

Single fly was crushed in 100 µL of extraction buffer (100 mM NaCl; 10 mM Tris-HCl, pH 8.0; 1 mM EDTA, pH 8.0; 2% Triton X-100; 1% SDS), and mixed with 100 µL of phenol/chloroform (1:1). After centrifugation, the aqueous layer was mixed with ethanol at 1:3 ratio, and centrifuged. Precipitated DNA was dissolved in TE buffer. Amplicon sequencing of 16S rDNA , which is targeted to 250 bp DNA encoding the bacterial 16S rRNA V4 region, was performed using the “early-pooling” method as previously described (*59*). The 1st PCR was performed with a 20-µL reaction volume containing 10.0 µL of 2 × Platinum SuperFi II Master Mix (Thermo Fisher Scientific, Waltham, MA, USA), 6.0 µL of each 1.66 µM 1st PCR F/R primer (515F/806R) (*59*), and 4.0 µl of the template DNA. The thermal cycle profile after an initial 30 sec denaturation at 98°C was as follows (35 cycles): denaturation at 98°C for 10 sec; annealing at 60°C for 10 sec; and extension at 72°C for 15 sec, with a final extension at the same temperature for 5 min. Each PCR product was purified using ExoSAP-IT Express (Thermo Fisher Scientific, Waltham, MA, USA). The purified PCR products were pooled and further purified, and the concentration was adjusted using AMPure XP (PCR product: AMPure XP beads = 1:0.8; Beckman Coulter, Brea, CA, USA), which was used as templates for the 2nd PCR. The 2nd PCR was performed to append Illumina adaptors with a 20-µL reaction volume containing 10 µL of 2 × Platinum SuperFi II PCR Master Mix, 8.0 µL of each 1.25 µM 2nd PCR F/R primer (*59*), and 2.0 µL of the pooled 1st PCR product. The thermal cycle profile after an initial 30 sec denaturation at 98°C was as follows (10 cycles): denaturation at 98°C for 10 sec; annealing/extension at 72°C for 15 sec, with a final extension at the same temperature for 5 min. The 2nd PCR product was purified using AMPure XP (PCR product: AMPure XP beads = 1:0.8). The target-sized DNA of the purified library was excised using E-Gel SizeSelect (ThermoFisher Scientific, Waltham, MA, USA). The DNA concentrations of the library was quantified using a Quantus Fluorometer (Promega, Madison, WI, USA). The DNA concentration of the library was adjusted, and then, sequenced on the NovaSeq system (2 × 250 bp PE; Illumina, San Diego, CA, USA).

### Bioinformatic analysis

To process RNA sequencing data, the quality of the sequencing reads was initially evaluated using FastQC (version 0.11.4) (http://www.bioinformatics.babraham.ac.uk/projects/fastqc/), and proceeded to adaptor trimming using Trimmomatic (*60*), followed by mapping to the *Drosophila* reference genome, dm6, from UCSC using STAR (*61*). Reads with mapping quality below eight and non-primary mapped reads were eliminated. The filtered reads were analyzed with HTSeq-count (*62*) to obtain the number of reads mapped on exons (1.75-3.68 million reads), which were further analyzed and normalized on R (*63*) using DESeq2 (*64*). PCA. The relative rpkm value was visualized and color-coded using Java TreeView (*65*). After obtaining the shared genes, STRING (*66*) was used to obtain the PPI (protein–protein interactions) with a 0.7 in confidence as minimum required interaction score. Cytoscape V3.9.1 software (*67*).

To determine the taxonomy of microbiome from 16S rRNA amplicon sequences, we processed the obtained data as previously described (*59*). Raw sequences were demultiplexed using cutadapt (*68*) and quality-filtered using fastp (*69*). Then, 515F/806R primer region was trimmed by cutadapt (*68*). The primer-trimmed sequences were processed using DADA2 (*70*), an amplicon sequence variant (ASV) approach. At the quality filtering process in DADA2, low quality and unexpectedly short reads were removed using DADA2::filterAndTrim() function with arguments of truncLen = c(210,200), and maxEE = c(2,2). Error rates were learned using DADA2::learnErrors() function. Then, sequences were dereplicated, error-corrected using DADA2::dada() with an argument pool = TRUE, and merged to produce an ASV-sample matrix. Chimeric sequences were removed using the DADA2::removeBimeraDenovo() function. Taxonomic identification was performed for ASVs based on the query-centric auto-k-nearest-neighbor (QCauto) method (*71*) and subsequent taxonomic assignment with the lowest common ancestor algorithm (*72*), implemented in Claident (https://www.claident.org/). The ASVs which were not assigned to a specific genus were separately analyzed using BLAST (*73*) to identify their genus.

### Quantification of transcripts (RT-qPCR)

To examine *SPH93* expression in flies fed with live bacteria, total RNA was extracted from one whole body using TRIzol reagent (15596026, Thermo Fisher Scientific, San Jose, CA, USA), and 125 μg of RNA was used to synthesize cDNA using ReverTra Ace qPCR RT Master Mix with gDNA Remover (FSQ-301, TOYOBO, Osaka, Japan). The cDNAs were then analyzed using quantitative real-time PCR (Bio-Rad Laboratories, Hercules, CA, USA).

To quantify the amount of 16S rDNA, groups of 10 flies were crushed in 400 µL of extraction buffer (100 mM NaCl; 10 mM Tris-HCl, pH 8.0; 1 mM EDTA, pH 8.0; 2% Triton X-100; 1% SDS), and mixed with 400 µL of phenol/chloroform (1:1). After centrifugation, the aqueous layer was mixed with ethanol at 1:3 ratio, and centrifuged. Precipitated DNA were dissolved in TE buffer and analyzed by real-time qPCR, using the primers amplifying the conserved region in 16S rDNA and fly genomic *Gapdh2* gene locus as follows: 16S rDNA, 5’-TCCTACGGGAGGCAGCAGT-3’, 3’-GGACTACCAGGGTATCTAATCCTGTT-5’; *Gapdh2*, 5’-CTGGCATTTCGCTAAACGAC-3’, 3’-CCTTGCTCTGCATGTACTTG-5’.

To quantify the amount of 16S rDNA specific to *A. junii*, groups of 10 flies were crushed in 100 µL TE buffer containing 0.1% SDS, and after centrifugation, the supernatant was removed. The supernatant contained fly genomic DNA, and approximately half of 16S rDNA is concentrated in the pellet which is recovered in 400 µL TE buffer containing 0.1% SDS with a brief sonication. After phenol/chloroform extraction, DNA was ethanol-precipitated. The DNA obtained was dissolved in 20 µL TE buffer, which was analyzed by real-time qPCR. Removing fly genomic DNA and extracts allows successful amplification of 16S rDNA from *A. jinii*. The primers amplifying the variable 5’ UTR region of 16S rDNA specific to *A. jinii* was used: 5’-GCAACCCTTCTGAAATGCTG-3’, 3’-AACCAACACAAGAATCTAACCAC-5’.

### Metabolite extraction and a widely targeted metabolomics

To determine the metabolites in fly heads, widely targeted metabolomics analysis was performed as previously described (*74, 75*). In brief, frozen samples in 1.5 mL plastic tubes were homogenized in 300 µL cold methanol with a f3 mm zirconia bead using a freeze crusher (TAITEC) at 41.6 Hz for 2 min. The homogenates were mixed with 200 µl of methanol, 200 µL of H_2_O, and 200 µL of CHCl_3_ and vortexed for 20 min at room temperature. The samples were then centrifuged at 15,000 rpm (20,000 g) for 15 min at 4℃. The insoluble pellets were used to quantify the total protein using a BCA protein assay kit (Thermo). The supernatant was mixed with 350 µL of H_2_O, vortexed for 10 min at room temperature, and centrifuged at 15,000 rpm for 15 min at 4℃. The aqueous phase was collected and dried down using a vacuum concentrator. The samples were re-dissolved in 2 mM ammonium bicarbonate (pH 8.0) containing 5% methanol and analyzed by LC-MS/MS. Chromatographic separations in an ACQUITY UPLC H-Class System (Waters) were carried out under reverse-phase conditions using an ACQUITY UPLC HSS T3 column (100 mm × 2.1 mm, 1.8 µm particles, Waters) and under HILIC conditions using an ACQUITY UPLC BEH Amide column (100 mm × 2.1 mm, 1.7 µm particles, Waters). Ionized compounds were detected using a Xevo TQD triple quadrupole mass spectrometer coupled with an electrospray ionization source (Waters). The peak area of a target metabolite was analyzed using MassLynx 4.1 software (Waters). The metabolite signals were normalized to the total protein level of the corresponding sample after subtracting the values from the blank sample.

The metabolites in the bacterial cultured media were determined via ion chromatography-high resolution mass spectrometry (IC-HR-MS) and LC-MS/MS as previously described (*76*). The media after removing the bacteria by centrifugation were sonicated in ice-cold methanol (500 μL), to which an equal volume of chloroform and 0.4 times the volume of ultrapure water (LC/MS grade, Wako Pure Chemical, Tokyo, Japan) were added. The suspension was centrifuged at 15,000 *× g* for 15 min at 4 °C. The aqueous phase was then filtered in an ultrafiltration tube (Ultrafree MC-PLHCC, Human Metabolome Technologies, Tsuruoka City, Japan), and the filtrate was concentrated using a vacuum concentrator (SpeedVac; Thermo Fisher Scientific, Waltham, MA, USA). The concentrated filtrate was dissolved in 50 μL ultrapure water and subjected to IC-HR-MS and LC-MS/MS analyses. Methionine sulfone (L-Met) and 2-morpholinoethanesulfonic acid (MES) were used as internal standards for cationic and anionic metabolites, respectively.

In IC-HR-MS, metabolites were detected using an orbitrap-type MS instrument (Q-Exactive focus; Thermo Fisher Scientific) connected to a high-performance IC system (ICS-5000 + , Thermo Fisher Scientific) that enabled highly selective and sensitive metabolite quantification owing to the IC separation and Fourier transfer MS principle (*77*). The IC instrument was equipped with an anion electrolytic suppressor (Dionex AERS 500; Thermo Fisher Scientific) to convert the potassium hydroxide gradient into pure water before the sample entered the MS instrument. Separation was performed using a Dionex IonPac AS11-HC 4 μm particle size column (Thermo Fisher Scientific). The IC flow rate was 0.25 mL/min, supplemented post-column with 0.18 mL/min makeup flow of MeOH. The potassium hydroxide gradient conditions for IC separation were as follows: from 1 mM to 100 mM (0-40 min) to 100 mM (40-50 min) and to 1 mM (50.1-60 min) at a column temperature of 30°C. The mass spectrometer was operated in the ESI-negative mode for all detections. A full mass scan (m/z 70–900) was performed at a resolution of 70,000. The automatic gain control target was set at 3 × 10^6^ ions, and the maximum ion injection time was 100 ms. The source ionization parameters were optimized with a spray voltage of 3 kV, and other parameters were as follows: transfer temperature, 320°C; S-Lens level, 50; heater temperature, 300°C; sheath gas, 36; and aux gas, 10.

Cationic metabolite concentrations were determined by liquid chromatography-tandem mass spectrometry (LC-MS/MS) as previously described (*76*). We employed a triple-quadrupole mass spectrometer equipped with an electrospray ionization (ESI) ion source (LCMS-8060, Shimadzu Corporation, Kyoto, Japan) operated in both positive and negative-ESI and in multiple reaction monitoring (MRM) modes. Analyte separation was achieved on a Discovery HS F5-3 column (2.1 mm I.D. × 150 mm L, 3 μm particle size; Sigma-Aldrich, St. Louis, MO, USA) through a gradient elution with mobile phase A (0.1% formate) and mobile phase B (acetonitrile containing 0.1% formate). The elution profile was as follows: 100:0 (0-5 min), 75:25 (5-11 min), 65:35 (11-15 min), 5:95 (15-20 min), and 100:0 (20-25 min), with a constant flow rate of 0.25 mL/min and a column oven set at 40°C.

For the measurement of metabolites registered in our in-house compound library, we compared the measurement results of samples and corresponding standards and confirmed that the retention times were consistent. Chromatographic peak integrations and confirmation of signal specificity for target compounds were performed for IC-HR-MS and LC-MS/MS, respectively, using Trace Finder software (ver. 4.1, Thermo Fisher Scientific) and Lab Solutions software (ver. 5.113, Shimadzu). For IC-HR-MS data analysis, the Trace Finder compound identification and confirmation setup parameters included a molecular ion intensity threshold override of 10,000, S/N 5, and mass tolerance of 5 ppm. Isotope pattern analysis using a 90% fit threshold, 30% allowable relative intensity deviation, and 5 ppm mass deviation were also performed to ensure that the relative intensities of the M + 1 and/or M + 2 isotope peaks for each compound were consistent with the theoretical relative intensities. For LC-MS/MS analysis, chromatographic peaks obtained with compound-specific SRM channels were integrated and manually reviewed. For a single target compound, one or more confirmatory SRM channels were set (if available) to confirm peak compound identification. Chromatograms were acquired using Lab Solutions (ver. 5.113, Shimadzu). Peak areas were determined using Data browser software. The obtained peak quantitation values for each compound were corrected for recovery by IS (MES and L-Met for IC-HR-MS and LC-MS/MS, respectively). For the tissue metabolome, peak area values were further corrected for weight.

### Statistical analysis

No statistical calculations were used to predetermine sample sizes. Our sample size was similar to those generally used in this field of research. Flies from each cross were randomly assigned to the treatment groups where possible. For behavioral experiments, all samples were numbered, and the investigators were blinded. For other experiments, blinding was not performed. Statistical analyses were performed using Prism version 10. For behavioral data analysis, the Mann-Whitney U test was used for comparisons between two groups, and the Kruskal-Wallis test followed by Dunn’s multiple comparison test was used for comparisons among multiple groups. *P* values < 0.05 were regarded as statistically significant. For boxplots, lower and upper whiskers represent 1.5 × inter quartile range (IQR) of the lower and upper quartiles, respectively, while boxes indicate upper quartile, median, and lower quartile from the top. In the graph with all data plots, data are presented as median ± IQR. Each experiment was successfully reproduced at least twice and performed on multiple days.

## Acknowledgments

We thank all lab members for their helpful discussions. We would like to thank Yi Rao, Ronald L Davis, Minoru Saitoe, Yan Yan, Bloomington Drosophila Stock Center (NIH P40OD018537), and Vienna Drosophila RNAi Center (VDRC, https://vdrc.at) for the materials. We also thank HKUST Biosciences Central Research Facility (BioCRF) team for their assistance.

## Funding

JST PRESTO program (YH)

The Fei Chi En Education and Research Fund (YH) Kato Memorial Bioscience Foundation (YH)

The Sumitomo Foundation (YH)

The Suzuken Memorial Foundation (YH) The Novartis Foundation (YH)

The Ichiro Kanehara Foundation (YH) Senri Life Science Foundation (YH)

The joint research program of the Institute for Molecular and Cellular Regulation, Gunma University (TN, YH).

The Big Data for BioIntelligence Laboratory (Z0428) from The Hong Kong University of Science and Technology (YL).

## Author contributions

Conceptualization: PB, YH

The fly experiments: PB, SC

Metabolome from flies: PB, YY, TN

Metabolome from bacteria: PB, YS

Metabolite feeding: PB, HN

Bacterial 16S rDNA sequencing: PB, MU

Analysis of bacterial load: PB, KFC

Analysis of spGFP flies: YSF, YH

Fat body analysis: PB, ME

Analysis of gut integrity: PB, HEJ, JK

Establishing transgenic bacteria: JL, YL

Supervision: PB, YL, YH

Writing – original draft: PB, YH

Writing – review & editing: PB, YY, MU, TN, KH, ME, JK, HEJ, YL, YH

## Data and materials availability

All data are available in the main text or the supplementary materials. The GEO accession number for the RNA-seq data and 16S rDNA amplicon sequencing reported in this paper are GSE294309, GSE294444, respectively. The code to analyze the data of 16S rDNA amplicon sequencing is deposited (https://github.com/ong8181/hiranolab_sleep-16SrRNA-amplicon).

