## supplemental info for "Microbiome-host interactions drive a trade-off between sleep quality and lifespan in *Drosophila*": biorxiv_suppl. materials_251224.pdf

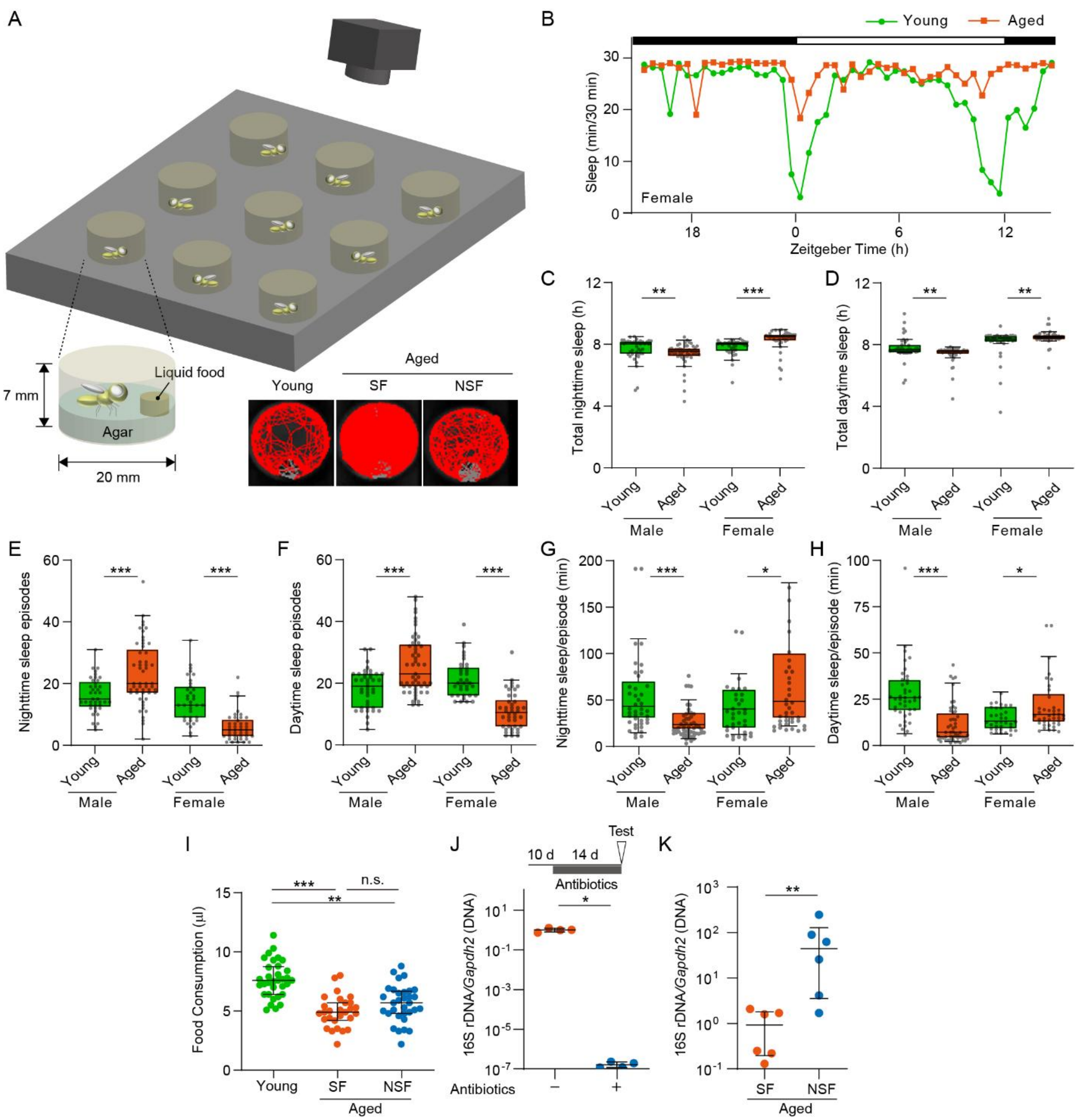


**Fig. S1. Characterization of the aged population showing SF or NSF.**

**(A)** Schematics of fly arena. A representative trace of fly locomotion is shown.

**(B)** Sleep profile of females. The 5-day-old and 25-day-old female flies were video-recorded to monitor the sleep duration every 30 min, and the averages were plotted against the Zeitgeber time (ZT). The light:dark cycle was indicated by white/black bars. Young, $n = 35$; aged, $n = 42$.

**(C and D)** Total sleep during nighttime between ZT13-23 (C) or daytime between ZT1-11 (D) of males and females. Two-sided Mann-Whitney U-test; from left, $n = 41, 53, 35, 42$.

**(E and F)** Sleep event number (episodes) during nighttime between ZT13-23 (E) or daytime between ZT1-11 (F) of males and females. Two-sided Mann-Whitney U-test; from left, $n = 41, 53, 35, 42$.

**(G and H)** Average nighttime or daytime sleep duration in each sleep episode. The total sleep duration was divided by the number of sleep episodes during nighttime between ZT13-23 (G) or daytime between ZT1-11 (H). Two-sided Mann-Whitney U-test; from left, $n = 41, 53, 35, 42$.

**(I)** Aged males with SF or NSF exhibit similar food intake. The food intake was estimated by measuring the reduction in liquid food after 36 h of video-recording. Kruskal-Wallis test, $P < 0.0001$; from left, $n = 33, 28, 32$.

**(J)** Feeding flies antibiotics reduces the bacterial load. Flies at 10-day old were fed antibiotics for 14 days and pooled. The bacterial 16S rDNA from the flies were analyzed via qPCR, which were normalized by fly genomic *Gapdh2*. Two-sided Mann-Whitney U-test; $n = 4$ for all.

**(K)** Aged males with SF exhibit lower bacterial load. Sleep was analyzed using aged males at 25-day old, and the flies with SF or NSF were pooled. The bacterial 16S rDNA were analyzed via qPCR, which were normalized by fly genomic *Gapdh2*. Two-sided Mann-Whitney U-test; $n = 6$ for all.

n.s., not significant $P > 0.05$; *, $P < 0.05$; **, $P < 0.01$; ***, $P < 0.001$.

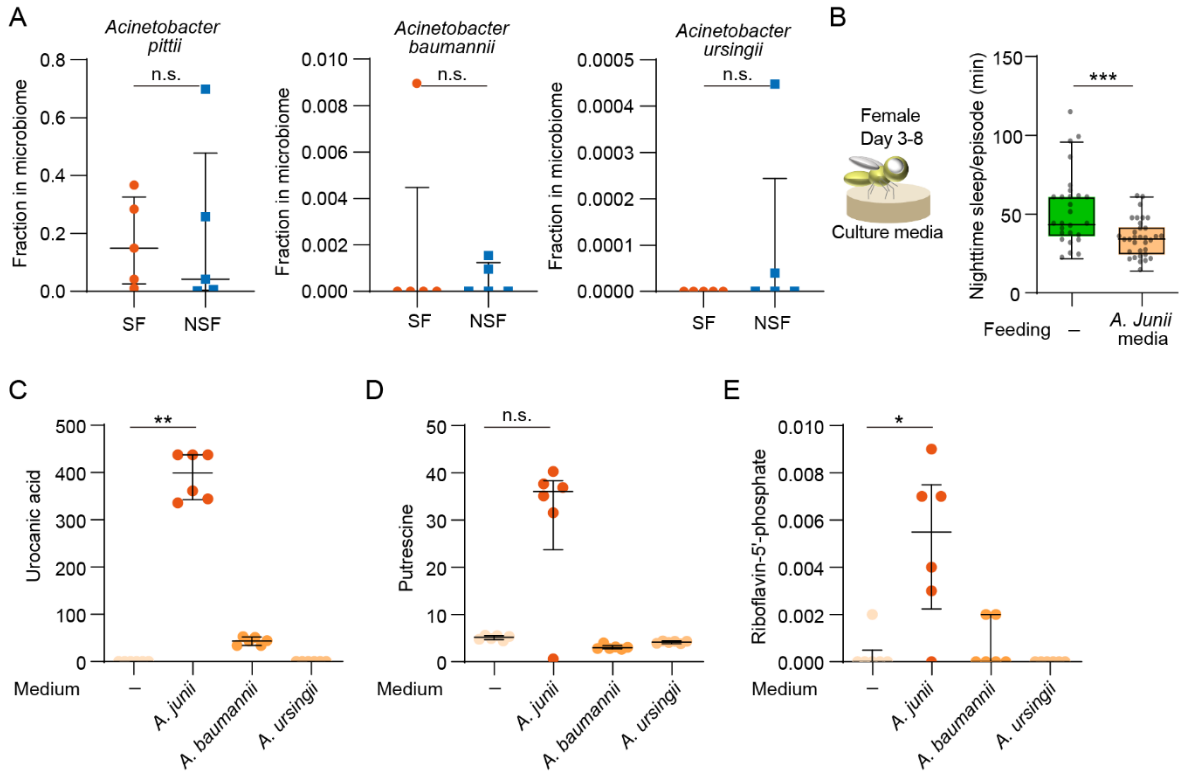


**Fig. S2. Metabolites enriched in the cultured media of *Acinetobacter junii*.**

**(A)** The 16S rDNA amplicon sequencing results were assigned to each indicated species, and each population of *Acinetobacter* was shown. Two-sided Mann-Whitney U-test; $n = 5$ for all.

**(B)** Feeding young females the culture media of *Acinetobacter junii* induces SF. Female flies at 3-day old were fed liquid food containing 80% of the media in which *A. junii* was cultured. Sleep was analyzed after feeding for 5 days. Kruskal-Wallis test, $P < 0.0001$; $n = 27, 33$.

**(C to E)** The abundance of metabolites in the bacterial culture media. Each *Acinetobacter* was cultured overnight and the media excluding the live bacteria was analyzed by LC-MS/MS. The medium without culturing bacteria was used as a control (medium -). Kruskal-Wallis test, (C) $P = 0.0002$, (D) $P =0.0033$, (E) $P = 0.0050$; $n = 6$ for all.

n.s., not significant $P > 0.05$; *, $P < 0.05$; **, $P < 0.01$; ***, $P < 0.001$.

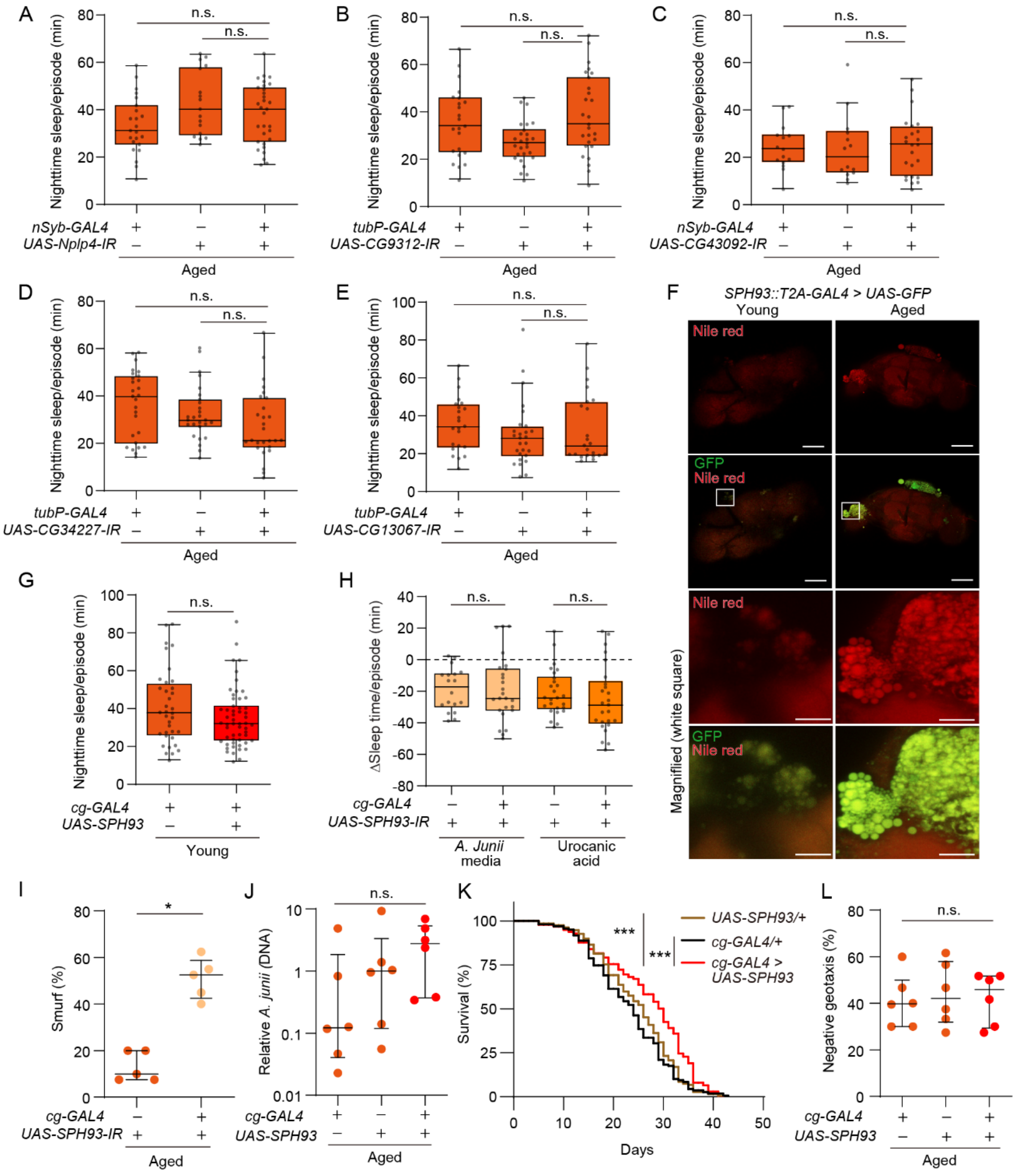


**Fig. S3.** ***SPH93*** **overexpression in fat body does not induce SF but extends lifespan.**

**(A to E)** Knockdown of type A genes enriched in aged SF male fly heads in Fig. 3D, other than *SPH93*, does not affect age-dependent SF. The indicated flies at 25-day old were examined by sleep assay. (A) Kruskal-Wallis test, $P = 0.1060$; from left, $n = 23, 17, 31$; (B) Kruskal-Wallis test, $P = 0.0210$; from left, $n = 23, 27, 27$; (C) Kruskal-Wallis test, $P = 0.8395$; from left, $n = 15, 14,$

24; (D) Kruskal-Wallis test, *P* = 0.0780; from left, *n* = 27, 27, 27; (E) Kruskal-Wallis test, *P* = 0.2639; from left, *n* = 23, 27, 23.

**(F)** *SPH93* is expressed in the fat body. The flies carrying the indicated transgenes at 5-day or 25-day old were fixed and the cuticles were removed, which was stained by Nile red to visualize the fat body. The images are representative of three experimental replicates. (upper) Scale bar: 100 □m. (lower) Scale bar: 20 □m.

**(G)** *SPH93* overexpression does not induce SF in young males. Sleep was analyzed using the indicated flies at 5-day old. Two-sided Mann-Whitney U-test; *n* = 36, 55.

**(H)** The reduced sleep duration by feeding *A. junii* and urocanic acid in *SPH93*-knockdown flies which was presented in Fig. 3G. Two-sided Mann-Whitney U-test; from left, *n* = 24, 18, 27, 25.

**(I)** Knockdown of *SPH93* increases leaky gut. The 25-day-old flies with the indicated transgenes were fed food containing blue dye for 1 day, and the flies showing blue dye in the body (Smurf flies) were counted. Two-sided Mann-Whitney U-test; *n* = 5 for all.

**(J)** Overexpression of *SPH93* does not change *A. junii* amount. The indicated flies at 25-day old were pooled and the abundance of *A. junii* was analyzed via qPCR. Kruskal-Wallis test, *P* = 0.1299; *n* = 6 for all.

**(K)** *SPH93* overexpression extends lifespan. Log-rank test, *n* = 250 for all.

**(L)** *SPH93* overexpression does not affect negative geotaxis in aged males. The indicated flies at 25-day old were tested by climbing assay. Kruskal-Wallis test, *P* < 0.9444; *n* = 6 for all.

n.s., not significant *P* > 0.05; *, *P* < 0.05; **, *P* < 0.01; ***, *P* < 0.001.

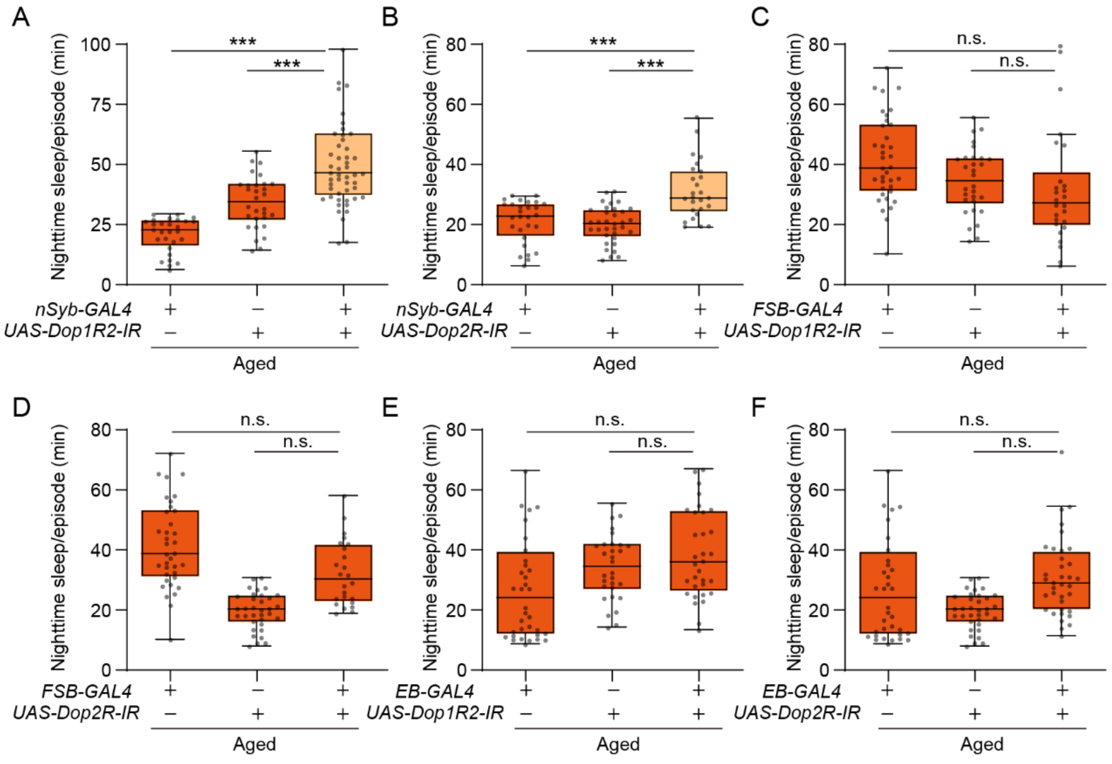


**Fig. S4. Involvement of *Dop1R2* and *DopR2* in age-dependent SF.**

**(A-F)** Neuronal knockdown of *Dop1R2* (A) and *DopR2* (B) suppresses age-dependent SF, but that in the FSB (C and D) or the EB (E and F) does not.

Sleep was analyzed using the indicated flies at 25-day old. *FSB-GAL4* and *EB-GAL4* used are *R75G12-GAL4* or *R15F02-GAL4*, respectively. (A) Kruskal-Wallis test, $P < 0.0001$; from left, $n = 28, 30, 52$. (B) Kruskal-Wallis test, $P < 0.0001$; from left, $n = 28, 34, 26$. (C) Kruskal-Wallis test, $P = 0.0067$; from left, $n = 36, 20, 36$. (D) Kruskal-Wallis test, $P < 0.0001$; from left, $n = 36, 24, 25$. (E) Kruskal-Wallis test, $P = 0.0117$; from left, $n = 36, 30, 32$. (F) Kruskal-Wallis test, $P = 0.0016$; from left, $n = 36, 34, 36$ .

n.s., not significant $P > 0.05$; *, $P < 0.05$; **, $P < 0.01$; ***, $P < 0.001$.

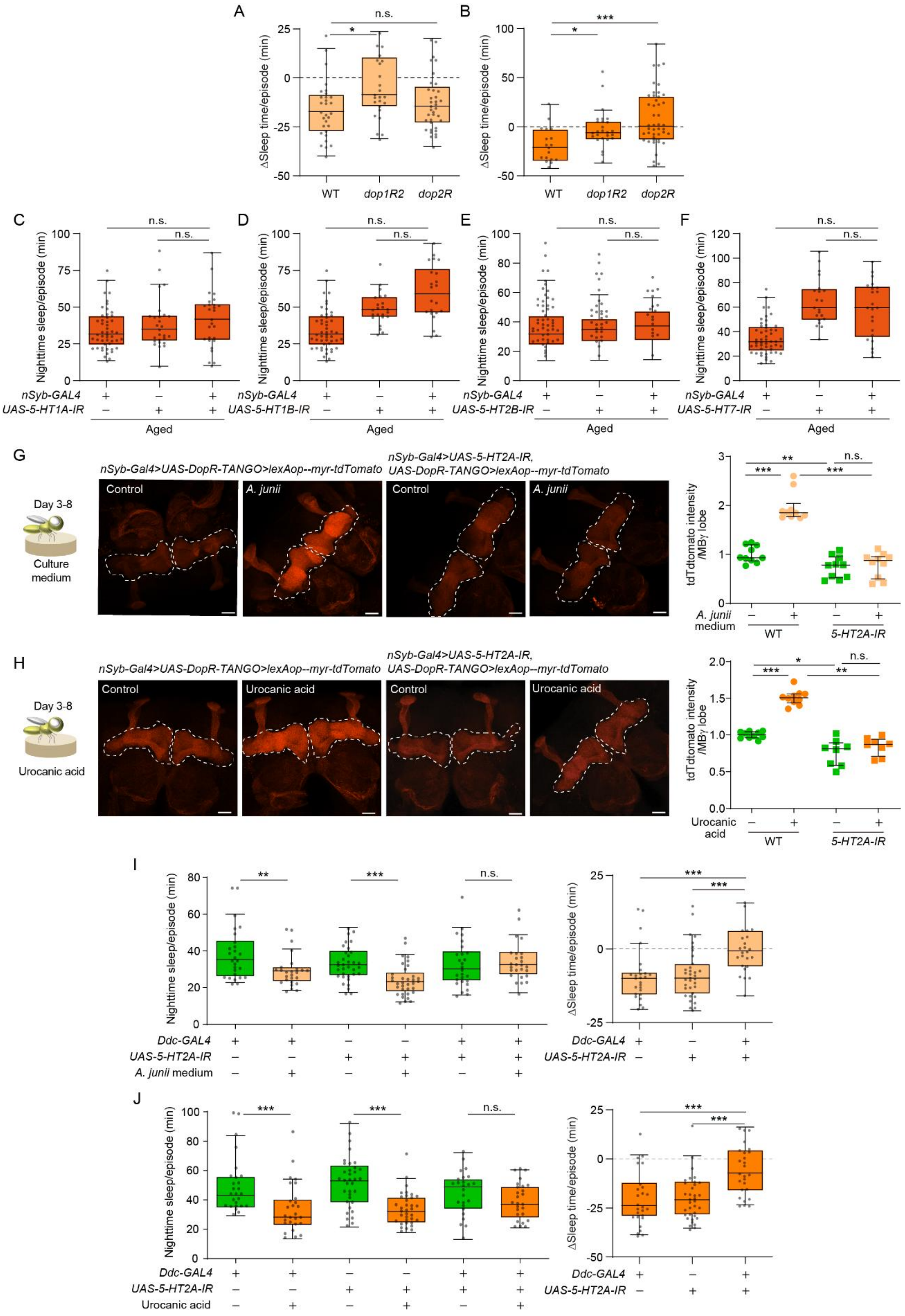
A
B
ΔSleep time/episode (min)
WT
dop1R2
dop2R
n.s.
*
***
C
D
E
F
Nighttime sleep/episode (min)
nSyb-GAL4
UAS-5-HT1A-IR
UAS-5-HT1B-IR
UAS-5-HT2B-IR
UAS-5-HT7-IR
Aged
G
Day 3-8
Culture medium
nSyb-Gal4>UAS-DopR-TANGO>lexAop--myr-tdTomato
nSyb-Gal4>UAS-5-HT2A-IR,
UAS-DopR-TANGO>lexAop--myr-tdTomato
Control
A. junii
tdTdtomato intensity /MBγ lobe
A. junii medium
WT
5-HT2A-IR
H
Urocanic acid
I
Ddc-GAL4
UAS-5-HT2A-IR
A. junii medium
J

**Fig. S5. Involvement of serotonin receptors in SF.**

**(A and B)** The reduced sleep duration by feeding *A. junii* (A) and urocanic acid (B) in *Dop1R2* and *DopR2* mutant flies, calculated from Fig. 5, B and C. (A) Kruskal-Wallis test, $P = 0.0186$; from left, $n = 30, 26, 38$. (B) Kruskal-Wallis test, $P = 0.0011$; from left, $n = 18, 26, 45$.

**(C to F)** Knockdown of serotonin receptors other than *5-HT2A* does not affect age-dependent SF. The indicated flies at 25-day old were examined by sleep assay. (C) Kruskal-Wallis test, $P = 0.2220$, from left, $n = 50, 26, 32$. (D) Kruskal-Wallis test, $P < 0.0001$; from left, $n = 50, 26, 32$. (E) Kruskal-Wallis test, $P < 0.0001$; from left, $n = 50, 26, 32$. (F) Kruskal-Wallis test, $P = 0.5735$; from left, $n = 50, 26, 32$.

**(G and H)** *5-HT2A* is required for upregulation of dopaminergic activity through feeding the culture media of *A. junii* (G) or urocanic acid (H). The 3-day-old flies with the indicated transgenes were fed food containing 50 mM urocanic acid or 80% of the *A. junii*-cultured medium for 5 days. The dissected brains were fixed and the fluorescence raw signal of tdTomato was quantified from MB g neurons (white dotted line). (G) Kruskal-Wallis test, $P < 0.0001$; $n = 10$ for all; (H) Kruskal-Wallis test, $P < 0.0001$; $n = 8$ for all.

**(I and J)** 5-HT2A in dopamine neurons is required for SF, induced by feeding *A. junii* media (I) or urocanic acid (J). Flies with the indicated transgenes at 3-day old were fed liquid food containing 80% of the *A. junii*-cultured medium (I) or 50 mM of urocanic acid (J) for 5 days and examined by sleep assay. The reduced sleep duration was presented on the right panel. (I, left) Kruskal-Wallis test, $P < 0.0001$; from left, $n = 27, 26, 34, 35, 27, 26$. (I, right) Kruskal-Wallis test, $P < 0.0001$; from left, $n = 26, 35, 26$. (J, left) Kruskal-Wallis test, $P < 0.0001$; from left, $n = 27, 27, 36, 35, 25, 26$. (J, right) Kruskal-Wallis test, $P < 0.0001$; from left, $n = 27, 35, 26$.

n.s., not significant $P > 0.05$; *, $P < 0.05$; **, $P < 0.01$; ***, $P < 0.001$.
